# Early post-infection treatment of SARS-CoV-2 infected macaques with human convalescent plasma with high neutralizing activity reduces lung inflammation

**DOI:** 10.1101/2021.09.01.458520

**Authors:** Koen K.A. Van Rompay, Katherine J. Olstad, Rebecca L. Sammak, Joseph Dutra, Jennifer K. Watanabe, Jodie L. Usachenko, Ramya Immareddy, Jamin W. Roh, Anil Verma, Yashavanth Shaan Lakshmanappa, Brian A. Schmidt, Clara Di Germanio, Nabeela Rizvi, Mars Stone, Graham Simmons, Larry J. Dumont, A. Mark Allen, Sarah Lockwood, Rachel E. Pollard, Rafael Ramiro de Assis, JoAnn L. Yee, Peter B. Nham, Amir Ardeshir, Jesse D. Deere, Jean Patterson, Aarti Jain, Philip L. Felgner, Smita S. Iyer, Dennis J. Hartigan-O’Connor, Michael P. Busch, J. Rachel Reader

## Abstract

Early in the SARS-CoV-2 pandemic, there was a high level of optimism based on observational studies and small controlled trials that treating hospitalized patients with convalescent plasma from COVID-19 survivors (CCP) would be an important immunotherapy. However, as more data from controlled trials became available, the results became disappointing, with at best moderate evidence of efficacy when CCP with high titers of neutralizing antibodies was used early in infection. To better understand the potential therapeutic efficacy of CCP, and to further validate SARS-CoV-2 infection of macaques as a reliable animal model for testing such strategies, we inoculated 12 adult rhesus macaques with SARS-CoV-2 by intratracheal and intranasal routes. One day later, 8 animals were infused with pooled human CCP with a high titer of neutralizing antibodies (RVPN NT_50_ value of 3,003), while 4 control animals received normal human plasma. Animals were monitored for 7 days. Animals treated with CCP had detectable levels of antiviral antibodies after infusion. In comparison to the control animals, they had similar levels of virus replication in the upper and lower respiratory tract, but had significantly reduced interstitial pneumonia, as measured by comprehensive lung histology. By highlighting strengths and weaknesses, data of this study can help to further optimize nonhuman primate models to provide proof-of-concept of intervention strategies, and guide the future use of convalescent plasma against SARS-CoV-2 and potentially other newly emerging respiratory viruses.

**Author summary:** The results of treating SARS-CoV-2 infected hospitalized patients with COVID-19 convalescent plasma (CCP), collected from survivors of natural infection, have been disappointing. The available data from various studies indicate at best moderate clinical benefits only when CCP with high titer of neutralizing antibodies was infused early in infection. The macaque model of SARS-CoV-2 infection can be useful to gain further insights in the value of CCP therapy. In this study, animals were infected with SARS-CoV-2 and the next day, were infused with pooled human convalescent plasma, selected to have a very high titer of neutralizing antibodies. While administration of CCP did not result in a detectable reduction in virus replication in the respiratory tract, it significantly reduced lung inflammation. These data, combined with the results of monoclonal antibody studies, emphasize the need to use products with high titers of neutralizing antibodies, and guide the future development of CCP-based therapies.

## INTRODUCTION

In late 2019, a newly identified coronavirus, SARS-CoV-2, began spreading rapidly across the globe [1]. While many infections are asymptomatic or result in mild symptoms, many people, especially those with predisposing factors, experience a severe acute respiratory syndrome (SARS), also called COVID-19, associated with > 1% mortality and often long-lasting complications.

In early 2020, the rapid surge in infection and hospitalization rates led to an urgent need for therapeutic interventions. This urgency sparked a high interest in collection and infusion of COVID-19 convalescent plasma (CCP), collected from people who had recovered from infection and had made antiviral antibodies, with particular focus on infusing CCP into recently infected patients with the hope of ameliorating their disease course. There were multiple rationales for the use of CCP. Despite variable evidence of efficacy, CCP therapy is a classic immunotherapy that has been applied to the prevention and treatment of many infectious diseases for more than a century, including more recently SARS, MERS, and the 2009 H1N1 influenza pandemic (reviewed in [2–4]). In addition, during a pandemic where many infected people survive and are willing to donate plasma, CCP can be collected by blood collection organizations (BCOs) and made readily available at relatively low cost. Finally, the reactivity of CCP evolves with the pandemic, as antibodies derived from recent convalescent survivors are expected to recognize recently circulating variants.

The early Emergency Use Authorization (EUA) by the Food and Drug Administration (FDA) of CCP for SARS-CoV-2 therapy in August 2020 was based on early promising results suggesting that the known and potential benefits of CCP outweighed any known and potential risks [5]. However, most of these early studies were observational, hindered by biases and other limitations. Since then, randomized clinical trials (including a meta-analysis) have indicated no therapeutic benefits of the average CCP, or at best, minimal benefits for CCP with high antiviral antibody titers given during the early stages of disease, prior to seroconversion including development of endogenous neutralizing antibodies [6–14]. Because of these later results, the FDA revised the EUA for CCP in February 2021, limiting the use to only high-titer CCP, and only in hospitalized patients who are early in the disease course or in those with impaired humoral immunity who cannot produce an adequate antibody response to control SARS-CoV-2 replication [15, 16].

Major hurdles for high-titer CCP-based strategies are that (i) most convalescent individuals don’t develop high titers of neutralizing antibodies [17, 18], (ii) the window of opportunity for CCP collection is limited, as neutralizing antibody titers decline over time [19, 20], and (iii) passive antibodies are unavoidably diluted in the recipient after CCP infusion. Thus, CCP-based therapies require careful screening of many plasma units to identify the relatively few donors who have a sufficiently high titer of antiviral antibodies. Currently the FDA defines high-titer CCP as having a neutralization titer of ≥250 (in the Broad Institute’s neutralizing antibody assay) or corresponding S/C cutoff thresholds defined by FDA for high-throughput binding antibody assays (e.g., ≥ 23 on the Ortho VITROS^®^ spike IgG assay, which was the first assay to be qualified for release of CCP by FDA) [15].

Considering the increased development and availability of potent neutralizing monoclonal antibodies, which despite high manufacturing costs, can be administered at high doses and have clearly proven efficacy [21, 22], it is unclear whether further investment in high-titer CCP-based strategies is scientifically and logistically merited, or what directions should be explored to make CCP more efficacious or cost-effective. For example, there is consideration of the potential collection and use of CCP derived from previously infected donors who were later vaccinated, which results in increased titers and breadth or neutralizing antibody reactivity against variants of concern (transfusion of CCP from vaccinated donors, including from vaccine-boosted previously infected donors, is not currently allowed according to the FDA EUA). Regardless, lessons gained from experience with SARS-CoV-2 CCP can be beneficial for rapid responses in future pandemics with other respiratory infectious agents.

Relevant animal models can be very helpful in understanding the efficacy of CCP and guiding this decision process. SARS-CoV-2 infection of nonhuman primates is a relevant animal model, because it recapitulates many of the key features of the human disease including high levels of virus replication, immunological responses to infection, and the development of interstitial pneumonia [23, 24]. The macaque model has been used to demonstrate the clear therapeutic efficacy of monoclonal antibodies [25–27]. Therapeutic studies with CCP in nonhuman primates have given mixed results. A pooled human CCP with moderate antibody titer given to rhesus macaques one day after virus inoculation failed to reduce virus replication (Deere et al, submitted for publication; [23]). In contrast, administration of a high-titer CCP, derived from convalescent African green monkeys, to African green monkeys 10 hours after inoculation had some therapeutic benefits, although variability and small group sizes limited statistical significance [28].

To further explore the potential benefit of CCP, and also to further validate the nonhuman primate model of SARS-CoV-2 to explore such passive immunotherapeutic interventions, the current study tested a pooled very high titer human CCP administered to rhesus macaques one day after high-dose virus inoculation, and compared it to animals treated with pooled control plasma. While administration of CCP did not result in a detectable reduction in virus replication in the respiratory tract, it significantly reduced lung inflammation.

## RESULTS

### Characterization of COVID convalescent plasma and control plasma

High-titer human CCP was prepared by mixing 3 plasma units (from 3 different donors), identified to have the highest titers of neutralizing antibodies (as determined by the reporter viral particle neutralization (RVPN) assay), and the highest reactivity of anti-spike total Ig (as determined by the Ortho VITROS^®^ IgG assay) [20]. The pooled CCP had NT_50_, NT_80_ and VITROS^®^ S/CO values of 3,003, 1,113 and 684, respectively, indicating high antiviral activity (**Fig. 1A, Table S1**). Similarly, control human plasma was prepared by mixing three control (i.e. pre-pandemic) plasma units that tested negative for neutralizing and binding antibodies using the same assays.

**Fig. 1.**
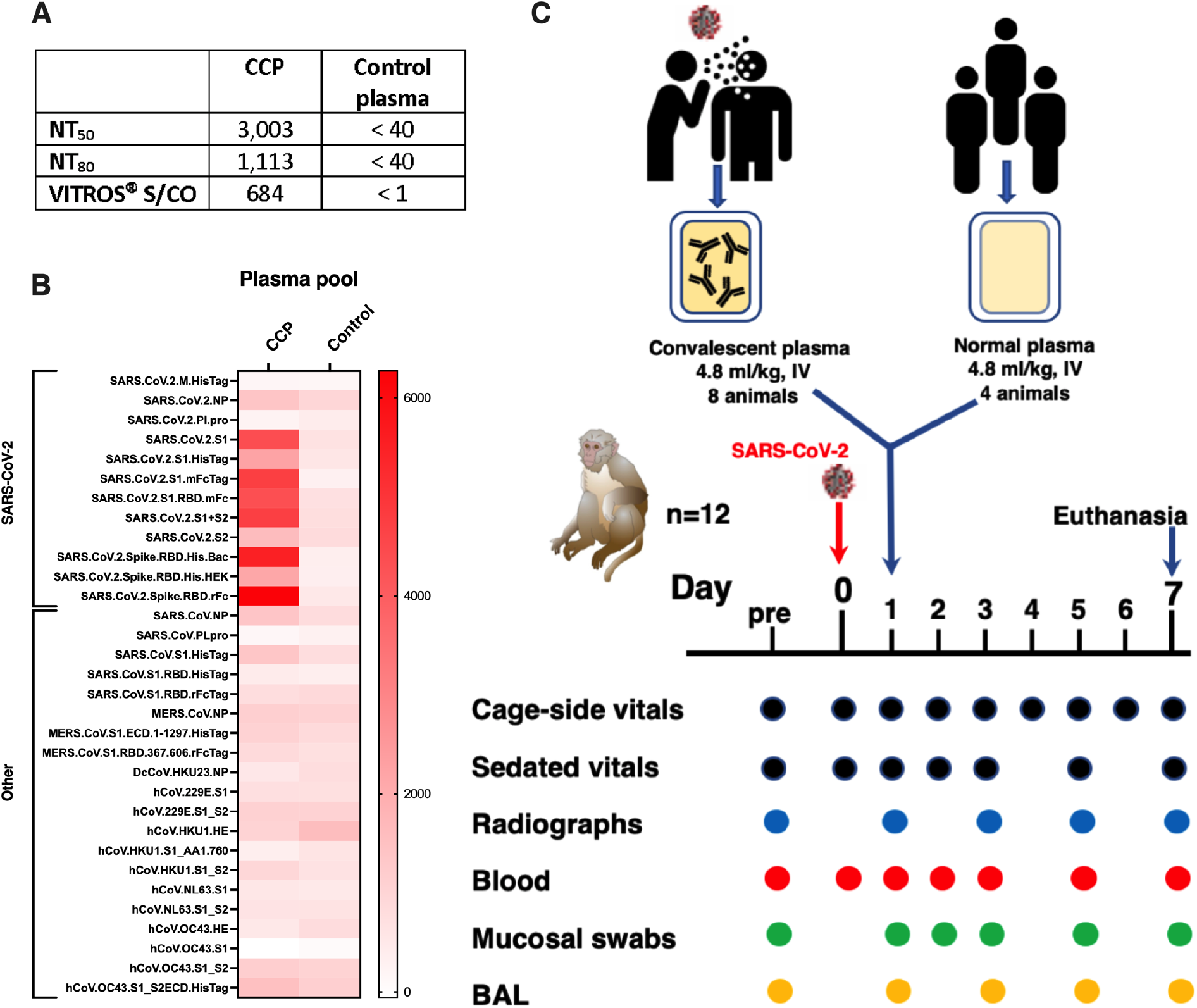
Experimental design & characterization of plasma pools. Two pools of human plasma, COVID convalescent plasma (CCP) and normal plasma, were prepared by mixing plasma of convalescent patients or pre-pandemic uninfected donors, respectively. (A) The 2 plasma pools were characterized for neutralizing antibody titers (NT50 and N80 values), and for total spike Ig (by VITROS^®^ assay. (B) The two plasma pools were also tested by coronavirus microarray assay (COVAM), and signal values are graphed as a heatmap. While the CCP had high reactivity to most SARS-CoV-2 antigens, cross-reactivity of the normal plasma pool to SARS-CoV-2 antigens was very low. Both plasma pools had similar reactivity to non-SARS-CoV-2 antigens. (C) Twelve adult rhesus macaques were inoculated on day 0 with SARS-CoV-2 by both intratracheal and intranasal routes. On day 1, eight animals received a single intravenous infusion with pooled CCP, while the other 4 animals received pooled normal control plasma. Animals were monitored closely for clinical signs (both cage-side and sedated observations) with regular collection of radiographs and samples to monitor infection and disease. On day 7, animals were euthanized for detailed tissue collection and analysis.

The pooled CCP and control plasma were also tested on a coronavirus antigen micro-array assay that detects antibodies against antigens of SARS-CoV-2, SARS-CoV-1, MERS and seasonal coronaviruses. As expected, the pooled CCP (but not the control plasma) had very high level reactivity against most SARS-CoV-2 antigens, while both CCP and normal plasma pools had similar low cross-reactivity to other coronaviruses (**Fig. 1B**).

### Experimental design to test therapeutic efficacy of COVID-19 convalescent plasma

Twelve young adult macaques were inoculated with a high dose (2.5 x10^6^ PFU) of a Washington isolate of SARS-CoV-2 by the intratracheal and intranasal routes. One day later, eight animals received a slow intravenous infusion with the pooled CCP (4.8 ml/kg), while the other four animals were treated with control plasma (4.8 ml/kg). Animals were monitored closely by clinical observation, radiographs, and regular sample collection, and were euthanized for tissue collection on day 7 (**Fig. 1C**).

### Detectable antiviral antibodies in serum after COVID-19 convalescent plasma administration

One day after infusion of plasma, all CCP-treated animals had detectable neutralizing activity in serum samples, as determined by RVNP and binding antibody assays (**Fig. 2**; **Table S2**). Between day 2 to 5, peak 50% neutralization titers (NT_50_) had a median value of 155, while NT_80_ values were near or below the assay’s limit of detection (titer of 40; Table S2). By day 7, some animals, including one control animal, had an increase in neutralizing activity in serum, indicative of a *de novo* antibody response. Similarly, starting one day after the infusion of CCP, all CCP-treated animals had detectable anti-spike immunoglobulins (as measured by the VITROS^®^ spike Total Ig assay), and reactivity to many SARS-CoV-2 antigens (as detected by the coronavirus antigen micro-array assay), which persisted throughout the observation period (**Table S2**; **Fig. 2B, Fig. S1**). Overall, the magnitude of the signals was as expected, based on the CCP being diluted ∼1:60-fold upon transfusion in the animals.

**Fig. 2.**
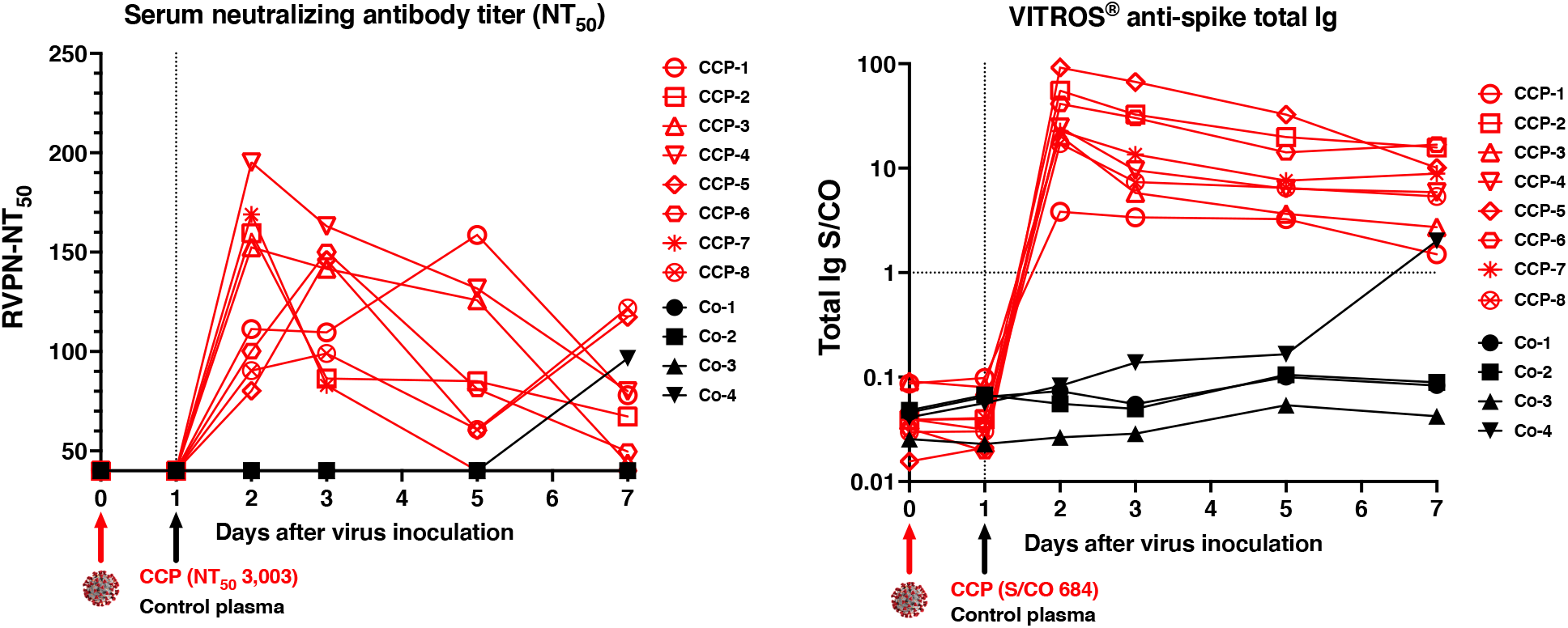
Neutralizing activity and anti-spike total Ig in serum of macaques after administration of convalescent or control plasma. Animals were inoculated on day 0 (red arrow) and administered either COVID convalescent plasma (CCP) or control (Co) plasma on day 1 (blue arrow). (A) Neutralizing activity was measured in serum samples of the animals using a RVPN assay, with estimation of the titer to get 50% inhibition (NT_50_). For comparison, the NT_50_ titer of the administered CCP was 3,003. Samples with undetectable titers are presented at the limit of detection (1:40). (B) VITROS^®^ anti-spike total Ig is expressed as the ratio of signal over cut-off (S/CO). A value of ≥1 indicates reactivity. The S/CO value of the administered pooled CCP was 684.

### Mild clinical disease irrespective of treatment

Animals were scored daily for several clinical parameters by cage-side observations. Overall, clinical signs were absent or mild-to-moderate (daily scores ≤ 4 out of a maximum of 22) and consisted mostly of nasal discharge. The highest daily score of 4 was recorded for CCP-treated animal 46174, which had a few observations of slightly increased abdominal breathing. When the sums of the daily scores from day 0 to 7 were tabulated for each animal, no significant difference was observed between the 2 treatment groups (**Fig. 3A-C**; p=0.28 Mann-Whitney).

**Fig. 3.**
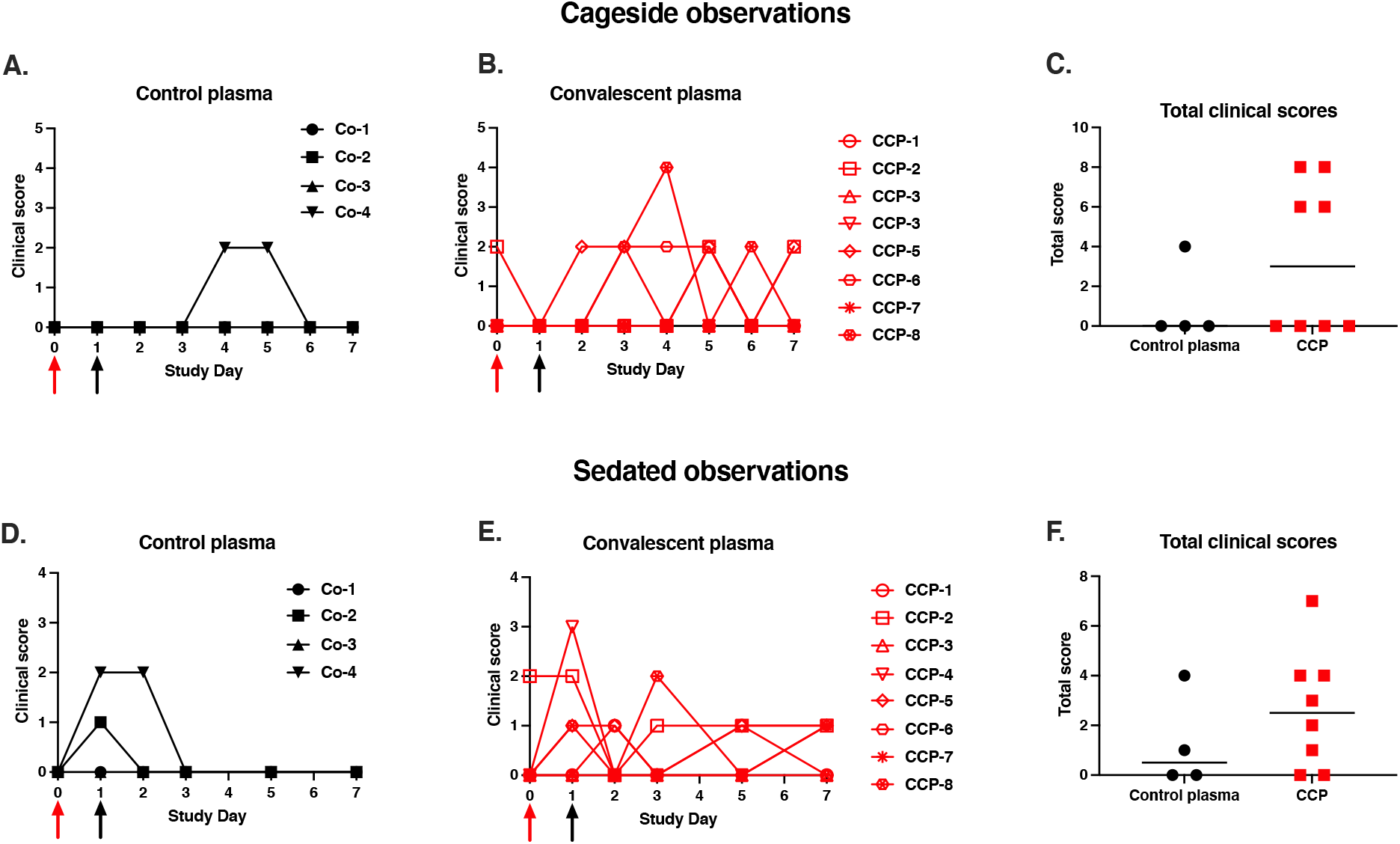
Mild clinical disease course with no detectable effect of convalescent plasma. Red and black arrows indicate time of virus inoculation and plasma administration on days 0 and 1, respectively. (A, B, D, E) Daily clinical scores based on cage-side observations and sedated measurements for each animal of the 2 study groups; the maximum daily score possible is 22 (for cage-side observations) and 27 (for sedated observations). (C, F) For each animal, the total of clinical scores over the 7-day period was tabulated. Comparison of the 2 groups revealed no detectable therapeutic benefits of the CCP treatment (p ≥ 0.28, Mann-Whitney).

Similarly, due to the mild disease course, there were no differences between the 2 groups for clinical scoring performed at time of sedation (**Fig. 3D-F**; p=0.41, Mann Whitney; **Fig. S2**). All animals had stable weights throughout the observation period, consistent with an adequate appetite. Three animals (2 CCP-treated animals and 1 control animal) had at least one recording of mildly elevated rectal temperature (102.5-103 ° F). None of the animals developed low oxygen saturation levels (Sp02 < 95%).

The lung radiological scores of animals in the study were normal (daily total score=0; n=6) to mild (daily total score 1-2; n=6) throughout the observation period, with no group differences (**Table S3**).

In summary, the overall mild clinical disease even in the control group, combined with the relatively small group sizes, made it difficult to use clinical markers and radiology as a measure of therapeutic efficacy of CCP.

### Innate and adaptive immune responses following infection

Following infection, most animals had a transient increase in C-reactive protein, ALT and AST, peaking on day 2, and a transient increase of several cytokines and chemokines, including IL-6, MCP-1, Eotaxin, I-TAC, IL-1RA and IP-10, generally peaking on day 1 or 2 (**Fig. S3-5**). Other cytokines and chemokines did not show consistent changes (**Fig. S4**). Overall, there were no discernible differences in these parameters between the 2 animal groups.

Analysis of immune subsets in whole blood revealed rapid dynamic changes in frequencies of innate immune cells as previously reported [23]. Relative frequencies of innate immune subsets in blood - neutrophils, proinflammatory monocytes, myeloid dendritic cells (mDC), plasmacytoid dendritic cells (pDC) - changed rapidly following infection in both experimental groups (**Fig. S6**). Assessment of T cell responses demonstrated a net decrease in naive CD4+ T cell frequencies at day 1 and day 7 post infection, while CD8+ T cell frequencies remained unchanged. Following infection, frequencies of both central memory (CD95+ CD28-) CD4+ and CD8+ T cells increased while effector memory (CD95+ CD28-) frequencies declined indicative of antigen driven activation and migration of T cells. In support of this, frequencies of Ki-67+ PD-1+ CD4 T cells were significantly increased at day 7 in both groups. Altogether, the data are consistent with infection-induced activation of the innate and adaptive arms in both the convalescent and normal plasma groups.

### COVID-19 convalescent plasma did not reduce virus replication in upper and lower respiratory tract

Nasal swabs, oropharyngeal swabs, and broncho-alveolar lavages (BAL) were collected regularly for viral load analysis. Samples were tested by RT-qPCR for total viral RNA (vRNA, N target), genomic viral RNA (gRNA, ORF1a target), subgenomic viral RNA (sgRNA, leader/N target), and cellular mRNA of the housekeeping gene PPIA (Peptidylprolyl Isomerase A). In general, and as previously shown [23, 27], the relative ratios of the 3 types of viral RNA’s were quite consistent in the samples (vRNA>gRNA>sgRNA).

sgRNA levels are considered the best evidence of active virus replication. Analysis of sgRNA in nasal and oropharyngeal swabs and BAL demonstrated that treatment with CCP at 1 day after infection did not have any detectable effect on virus replication in the upper or lower respiratory tracts, (**Fig. 4; Fig S7-8**). The data on total RNA and gRNA levels gave similar conclusions (**Fig. S7**).

**Fig. 4.**
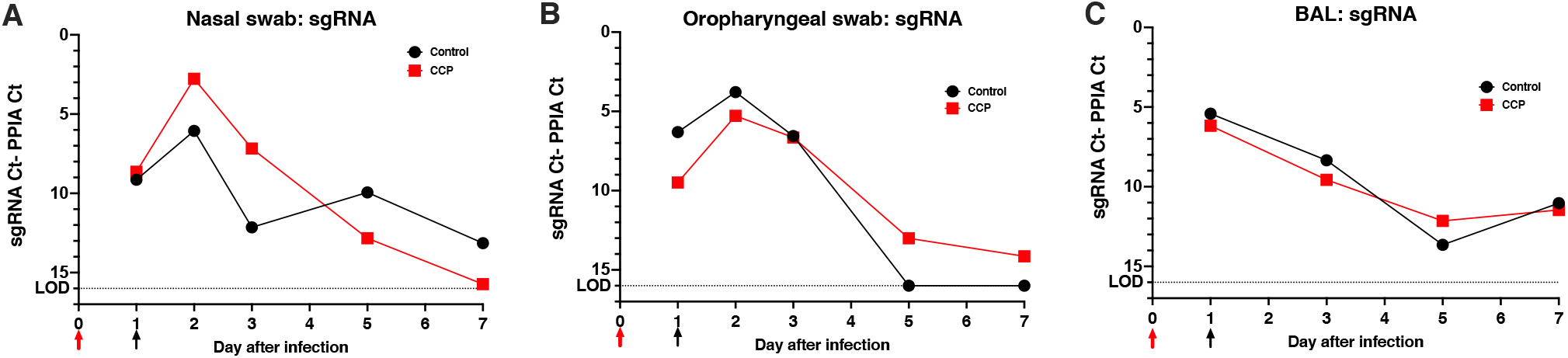
Lack of detectable effect of convalescent plasma on virus replication in upper and lower respiratory tract. (A) Time course of median viral subgenomic (sg) RNA copies in nasal swabs. (B) Time course of median viral sgRNA copies in oropharyngeal swabs. (C) Time course of median viral sgRNA in BAL samples. Red and black arrows indicate time of virus inoculation and infusion of plasma (control plasma or CCP) on days 0 and 1, respectively. sgRNA was measured by RT-qPCR and expressed relative to cellular mRNA of the housekeeping gene PPIA (as indicator of the cellular content in the sample tested) by plotting the difference in CT values; thus, a larger difference indicates less virus replication. The dotted line indicates the limit of detection (LOD). More details are provided in **Fig. S7**.

### COVID convalescent plasma treatment reduced lung inflammation

To evaluate infection-induced lung pathology, a comprehensive histology scoring system, described in detail in the methods and validated in an earlier study [27], was used to tabulate interstitial cellularity scores. This scoring system takes into consideration (i) interstitial pneumonia as the most striking and consistent lesion in the lungs of SARS-CoV-2 infected macaques at 7 days of infection, (ii) the multifocal to locally extensive highly random distribution of the lesions and absence of distinct borders, (iii) the requirement of x40 magnification for accurate evaluation of the severity of the lesions. An average of 1208 microscopic fields (range: 710-1634) per animal, representing all 7 lung lobes, were graded from 0 to 4 in a blind analysis to tabulate an overall interstitial cellularity score.

The CCP-treated animals had on average, a 17% lower interstitial cellularity score than the control plasma group, which was statistically significant (**Fig. 5**; p=0.006, t test). Thus, administration of CCP was associated with reduced lung inflammation 7 days after infection.

**Fig. 5.**
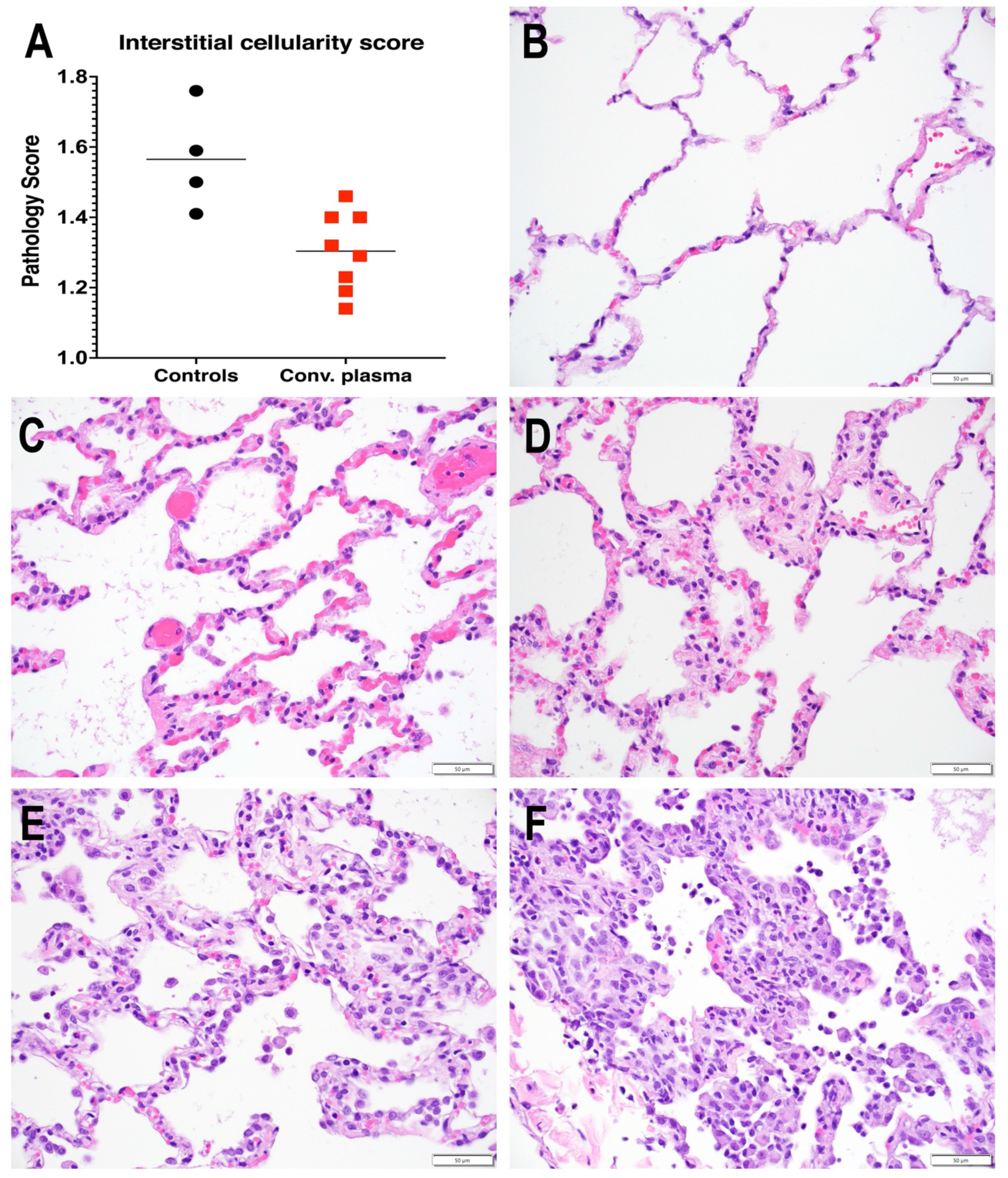
Reduced interstitial pneumonia in convalescent plasma-treated animals. **A.** Interstitial cellularity was evaluated on 7 lung lobes and an average score was tabulated as outlined in the methods section. Lines indicated mean values. The CCP group had significantly lower scores than the control group (p=0.006; unpaired t-test). **B-F.** Interstitial cellularity score assigned to random x40 fields is based on the number of cells expanding the alveolar interstitium. Representative x 40 images are shown. **B.** Grade 0: normal lung with thin acellular alveolar septae (animal CCP-7). **C.** Grade 1: alveolar interstitium expanded by 1 to 2 cells (animal CCP-1). **D.** Grade 2: alveolar interstitium expanded by 2 to 4 cells (animal CCP-1) **E.** Grade 3: alveolar interstitium expanded by 4 to 6 cells. (animal Co-3) **F.** Grade 4: alveolar interstitium expanded by more than 6 cells (animal Co-3)

### Multivariable analysis of correlates of efficacy

A multivariable analysis with correlation matrix was performed on the main data sets, including neutralization activity, virus replication, lung pathology scores and clinical scores. With the caveat of the small group sizes, lung pathology scores had a highly significant inverse correlation with peak serum NT_50_ values (Spearman r = −0.87; p < 0.001), followed by peak plasma VITROS anti-spike total Ig levels (Spearman r= −0.59, p = 0.046). In contrast, sgRNA viral loads in upper and lower respiratory tract secretions correlated poorly with lung pathology scores (**Fig. 6**). When only the data of the CCP-treated animals were analyzed, the negative correlation between NT_50_ peak titers and lung pathology scores was slightly reduced (Spearman r=-0.71, p=0.059), while NT_50_ peak titers correlated negatively with cage-side clinical scores (Spearman r=-0.77, p=0.04); **Fig. S9**.

**Fig 6.**
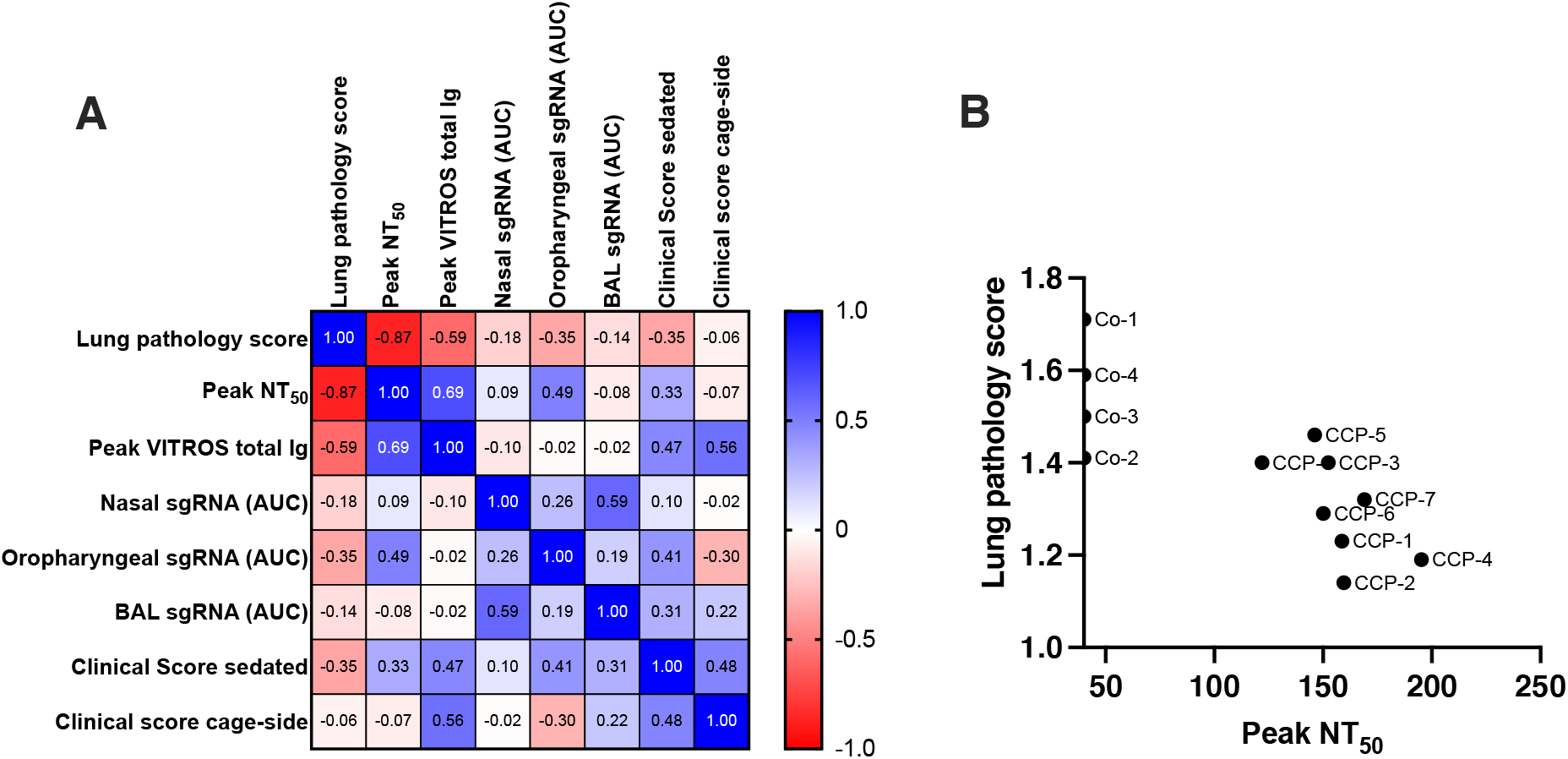
Multivariable correlation analysis. (A). Spearman r correlation matrix in heatmap format. For this analysis, lung pathology scores are the interstitial cellularity scores (from **Fig. 5**). Peak NT_50_ represents the peak neutralizing antibody titers up to day 5 (i.e., prior to possible *de novo* antibody responses). VITROS^®^ anti-spike total Ig represents the peak value for each animal (i.e., day 2; **Fig. 2**). Nasal, oropharyngeal and BAL sgRNA values are based on AUC of the data in **Fig. S7**. Clinical scores (sedated and cage-side) are the tabulated scores of each animal over the 7-day observation period (**Fig. 3C, F**). (B) Correlation between neutralizing antibody peak NT_50_ values and lung pathology scores (Spearman r = −0.87; p < 0.0005). The labels next to each symbol indicate the individual animals.

## DISCUSSION

The current study provides insights on the efficacy and limitations of CCP therapy against SARS-CoV-2 replication and COVID-19 disease, as well as the opportunities and challenges associated with the use of a nonhuman primate model in testing passive immunotherapy strategies.

We demonstrate that administration of a pooled human CCP with high titer of neutralizing and spike-binding antibodies, administered one day after virus inoculation, conferred therapeutic benefits to SARS-CoV-2 infected macaques in terms of reduced interstitial pneumonia, despite no detectable effect on reducing virus replication.

The lack of a detectable effect of the CCP on RNA levels in mucosal samples is likely multifactorial, with insufficient antiviral activity as the primary explanation, but influenced by additional experimental factors. Although we used a CCP with high neutralizing activity *in vitro*, the antibodies became diluted so much upon transfusion that by the time they reached mucosal sites, their concentration was probably too low to have a drastic impact on reducing virus replication *in vivo*. In this context, it is important to note that we inoculated animals with a very high dose of SARS-CoV-2, to induce rapid wide-spread infection of upper and respiratory tract, with peak virus replication occurring within the first 1-2 days. Having high levels of viral replication at the time of CCP administration sets a high bar to detect efficacy, especially as it takes time for passively infused antibodies to distribute and reach peak concentrations at mucosal sites. The difficulty to detect a difference was likely exacerbated by the considerable variability in virus levels in mucosal samples of SARS-CoV-2 infected animals, including untreated control animals, as observed in many other studies [25, 29–31]. This high variability is likely a combination of individual variability in virus replication, but also the variability inherently associated with mucosal secretion samples, which represent a snapshot in time of viral shedding at a limited mucosal surface. Thus, while small animal group sizes can still allow detection of large differences in virus replication caused by very potent antiviral strategies including passive immunotherapy with monoclonal antibodies, they lack the power to detect relatively mild-to-moderate antiviral effects.

In contrast to the lack of a detectable effect on virus replication, CCP treatment had a modest but statistically significant beneficial effect on reducing lung inflammation. The reason that the relatively modest difference (∼17%) in interstitial cellularity scores between the study groups was statistically significant, despite the relatively small group sizes, can be attributed to the very comprehensive and extensive scoring system, in which the evaluation of numerous microscopic fields per animal provides a relatively precise assessment of the overall extent of interstitial pneumonia at 7 days of infection. Despite its advantages of being highly rigorous and robust, this scoring system has the drawback of being labor-intensive, which precludes application on large-scale studies. Thus, future efforts can focus on further refining it, by simplification (i.e., pathologist-driven scoring of fewer fields or fewer lobes but achieving statistically similar reliable results) and/or automation (i.e., computer-generated scores).

There is precedence for a relative dissociation between SARS-CoV-2 virus replication, particularly in the upper respiratory tract, and pulmonary lesions, as demonstrated in several therapeutic studies in SARS-CoV-2 infected macaques [31, 32]. Dose-range vaccine studies in macaques found that higher antibody levels were needed to reduce virus replication in the upper airways than in the lower respiratory tract [33]; this can explain recent observations that some vaccinated people with breakthrough infections with the SARS-CoV-2 delta variant can have similar viral loads in upper respiratory tract as unvaccinated people, but yet, remain at much reduced risk for severe illness and hospitalization [34, 35]. Finally, it has been demonstrated in the lungs of SARS-COV-2 infected macaques that the virus does not necessarily co-localize with the lesions and can be found in areas of the pulmonary parenchyma that are not inflamed [36]. Altogether these observations underscore the importance of pulmonary histopathology as a key endpoint when evaluating the efficacy of therapeutics.

The data of the current study help to further define neutralizing activity as a correlate of efficacy for antibody-based antiviral therapeutic strategies against SARS-CoV-2. A previous study used the same experimental design as the current study, except that one day after virus, instead of CCP, animals received a combination of 2 potent monoclonal antibodies [27]. In that study, neutralizing antibody titers in serum after infusion were ∼ 2 to 3 logs higher than those observed in the current study. Despite similarly small group sizes, the antibody-treated animals had statistically significant reductions in virus replication, clinical signs, and interstitial pneumonia (∼ 50% reduction in interstitial cellularity scores) compared to control antibody-treated animals. Comparison of these 2 studies, with the lung histology scoring performed by the same pathologists, revealed that animals treated with monoclonal antibodies had significantly lower lung pathology scores than the CCP-treated animals in the current study (**Fig. S10**), indicating the superiority of monoclonal antibodies above CCP to treat early SARS-CoV-2 infection.

In the current CCP study, peak neutralizing antibody NT_50_ values in serum the first few days after infusion were ∼150, which, considering the marginal efficacy observed in the histology scoring, helps to set a threshold for neutralizing antibody titers in the recipient to have a chance at any therapeutic benefits. Although direct comparison of neutralization data across studies is difficult due to differences in assays and other laboratory conditions, the threshold value observed in this current study is consistent with recent findings from other animal and human studies. A study that used purified IgG derived from convalescent macaques found that a NT_50_ titer between 50 and 500 in the recipient animals was the threshold to see an effect on reducing virus replication, although no lung histology scoring was reported [37]. In a study with African green monkeys, the administration of a high-titer CCP, derived from convalescent African green monkeys, administered 10 hours after a moderate-dose virus inoculation, resulted in live-virus plaque reduction neutralization titers 50% (PRNT_50_) of ∼ 128 in the recipient animals, that were associated with reduced virus replication and histology, although differences were statistically not significant, likely due to variability and small group sizes [28]. In human studies, it has been difficult to set a threshold for neutralizing activity after CCP infusion, as generally infusions were performed later in the course of infection (i.e. when a *de novo* antibody responses were already being generated), or studies did not report neutralization titers in the CCP recipients [8, 10].

The combined results of these studies provide further guidance to what future, if any, CCP can have in the clinic for early treatment of SARS-CoV-2 infected people. First of all, as very few people who recover from COVID-19 develop persistently high neutralizing antibody titers, the use of CCP from such donors faces increasing logistical and scientific hurdles, especially considering the increased availability of potent monoclonal antibodies. However, recent studies have demonstrated that COVID-19 survivors who subsequently received a SARS-CoV-2 vaccine made very strong booster immune responses, which also neutralize currently circulating variants of concern [38–44]. Thus, vaccinated COVID-19 recovered subjects are likely to be a much better source of CCP.

The current study also helps to further validate and strengthen the nonhuman primate model of SARS-CoV-2 infection and COVID-19. Although the typical disease course of SARS-CoV-2 infection in young, otherwise healthy macaques is mild, and despite a limited number of animals, a detailed analysis was able to detect relatively mild-to-moderate therapeutic benefits of high-titer CCP administration. These findings will be relevant for future pandemics with newly emerging respiratory viruses, as the rapid development of relevant nonhuman primate models with proper monitoring and scoring systems can speed up testing the safety and efficacy of antiviral strategies including CCP and monoclonal antibodies to generate the data that can guide the design of clinical trials.

## Materials and methods

### Ethics Statement

The study was approved by the Institutional Animal Care and Use Committee of the University of California, Davis (study protocol 21735).

### Animals and care

All 12 rhesus macaques (Macaca mulatta) in the study were young adults (3.5 to 6.5 years of age), born and raised in the breeding colony of the California National Primate Research Center (CNPRC), which is negative for type D retrovirus, SIV and simian lymphocyte tropic virus type 1. Each of the 2 study groups had equal sex distribution (half males, half females) and similar age and weight (table S4). Prior to enrollment, animals were confirmed to be seronegative and RT-PCR negative for SARS-CoV-2 and were kept in a special barrier room prior to study initiation. Animals were moved into the animal biosafety level 3 (ABSL-3) facility just before virus inoculation.

The CNPRC is accredited by the Association for Assessment and Accreditation of Laboratory Animal Care International (AAALAC). Animal care was performed in compliance with the 2011 *Guide for the Care and Use of Laboratory Animals* provided by the Institute for Laboratory Animal Research [45]. Macaques were housed indoor in stainless steel cages (Lab Product, Inc.) whose sizing was scaled to the size of each animal, as per national standards, and were exposed to a 12-hour light/dark cycle, 64-84°F, and 30-70% room humidity. Animals had free access to water and received commercial chow (high protein diet, Ralston Purina Co.) and fresh produce supplements.

### Virus and inoculations

A virus stock of a Washington isolate was obtained from BEI Resources (SARS-CoV-2 2019-nCoV/USA-WA1/2020; NR-52352; Lot/Batch # 70033952). The titer of this stock was 10^6^ PFU/ml. Animals were inoculated with a total of 2.5 ml (2.5×10^6^ PFU), of which 2 ml was administered intratracheally via a 8 fr PVC feeding tube, and 0.5 ml was administered intranasally (0.25 ml per nostril).

### Convalescent and control plasma preparation and administration

Three convalescent plasma units, collected from 3 different patients who survived SARS-CoV-2 infection, and identified to have a high titer of neutralizing antibodies were selected for making a pool of human convalescent plasma. Considering the volume of human plasma that can be safely given to an animal, the weight of the 8 animals, and the limited amount available of the plasma with the highest titer, (NT80 of 2313), we decided to use a maximal amount of this highest-titer plasma and mix it with the 2 other high titer units at a ratio of 60:20:20 in order to administer the maximum absolute amount of convalescent plasma-derived neutralizing antibodies to the 8 animals (Table S1). A pool of control plasma was prepared by mixing three 2019 control (i.e., collected in 2019 prior to the SARS-CoV-2 pandemic) plasma units in equal amounts. Convalescent plasma and control plasma were administered to the animals at a dose of 4.8 mg/kg body weight, via slow intravenous infusion, approximately 24 hours after virus inoculation.

### Clinical observations and sample collections

Daily cage-side clinical monitoring was performed by a veterinarian who was blinded to the group assignments, and included recording of responsiveness, discharge, respiratory rate and character, evidence of coughing/sneezing, appetite, stool quality. A score was tabulated for each of these parameters, and a total score was calculated for each animal per day. When animals had to be sedated for procedures, additional clinical assessments (including rectal temperature, respiration, spO2, heart rate, skin turgor/hydration) were recorded by the same veterinarian. Details of the scoring criteria were published earlier [27]. Animals were sedated with ketamine HCl (10 mg/kg IM) for the clinical assessment. Dexmedetomidine (15 mcg/kg IM) was administered after clinical assessments to facilitate sampling, and midazolam (0.25-0.5 mg/kg IM) was added as needed. Oxygen saturation was obtained by pulse oximetry with a Radical 7 (Masimo, Irvine, CA). Blood pressure was obtained via oscillometry with a Vet25 and an appropriately sized cuff according to the manufacturer’s instructions (SunTech, Morrisville, NC).

Blood was collected via peripheral venipuncture. Complete blood counts were performed on EDTA-anticoagulated blood samples, with electronic cell counts performed on a Pentra 60C+ analyzer (ABX Diagnostics) or Vet abc™ (SCIL Animal Care); differential cell counts were determined manually. EDTA anti-coagulated blood was also used for immunophenotyping and, after centrifugation, the collection of plasma. Blood tubes without coagulant and CPT™ vacutainer tubes were also collected for processing via centrifugation (900xg for 10 minutes) for serum and peripheral blood mononuclear cells, respectively. Plasma and serum aliquots were stored at −70 °C until further processing.

Nasopharyngeal and oropharyngeal secretions were collected with FLOQSwabs™ (Copan), placed in a vial with DNA/RNA Shield™ solution (Zymo Research), and stored at −70 °C until further processing.

Bronchoalveolar lavages (BAL) were performed using a 20F rubber feeding tube with instillation of 20 ml sterile physiologic saline followed by aspiration with a syringe. BAL samples were spun in the lab. The BAL cell pellet, together with 0.5 ml of supernatant, was then mixed with 1.5 ml of TRIzol^®^-LS (Thermo Fisher Scientific) and cryopreserved at −70° C. Additional aliquots of BAL supernatant were also immediately cryopreserved.

At the end of the study, animals were euthanized, and a full necropsy was performed for tissue collection, including fixed tissues for histopathology (see further).

### Collection and evaluation of radiographs

Radiographs were obtained with a HF100+ Ultralight imaging unit (MinXRay, Northbrook, IL) at 50 kVp, 40mA, and 0.1 sec. Ventrodorsal, dorsoventral, R lateral, and L lateral radiographs were obtained prior to inoculation and every other day after virus inoculation (days 1, 3, 5, and 7). Radiographs were scored for the presence of pulmonary infiltrates by a board-certified veterinary radiologist, who was blinded to the experimental group and time point, according to a standard scoring system (0: normal; 1: mild interstitial pulmonary infiltrates; 2: moderate pulmonary infiltrates perhaps with partial cardiac border effacement and small areas of pulmonary consolidation; 3: severe interstitial infiltrates, large areas of pulmonary consolidation, alveolar patterns and air bronchograms). Individual lobes were scored and scores per animal per day were totaled.

### Viral load determination by RT-qPCR analysis

Quantitative real-time PCR assays were developed for detection of full-length genomic vRNA (gRNA), sub-genomic vRNA (sgRNA), and total vRNA. RNA was extracted from swabs preserved in DNA/RNA Shield using the Quick-RNA Viral Kit (Zymo Research). BAL cell pellets were processed directly in TRIzol-LS reagent (Thermo Fisher Scientific) and total RNA purified using the Qiagen RNeasy Mini Kit (Qiagen). Tissues preserved in RNA*later* (Sigma-Aldrich) were transferred to QIAzol (Qiagen), and homogenized with a 7mm stainless steel bead in a TissueLyser (Qiagen), and processed using the Qiagen RNeasy Mini Kit. Following DNase treatment with ezDNase (Thermo Fisher Scientific), complementary DNA was generated using random hexamers, Superscript IV Reverse Transcriptase (Thermo Fisher Scientific) in the presence of RNaseOUT (ThermoFisher). A portion of this reaction was mixed with QuantiTect Probe PCR Kit (Qiagen) and optimized concentrations of gene specific primers. All reactions were run on a Quantstudio 12K Flex real-time cycler (Applied Biosystems). gRNA was quantified by targeting orf1a-nsp4 using primers orf1a_F7 (GTGCTCATGGATGGCTCTATTA) and orf1a_R7 (CGTGCCTACAGTACTCAGAATC), with probe orf1a_P7 (/56-FAM/ACCTACCTT/ZEN/GAAGGTTCTGTTAGAGTGGT/3IABkFQ/). sgRNA was quantified using primers sgLeadSARSCoV2_F (CGATCTCTTGTAGATCTGTTCTC) and wtN_R4 (GGTGAACCAAGACGCAGTAT), with probe wtN_P4 (/56-FAM/TAACCAGAA/ZEN/TGGAGAACGCAGTGGG/3IABkFQ/). Total vRNA was quantified using primers wtN_F4 (GTTTGGTGGACCCTCAGATT) and wtN_R4, with probe wtN_P4. Standard curves generated from PCR amplicons of the qPCR targets were used to establish line equations to determine RNA copies/mL or copies/ug RNA. A *macaca* housekeeping gene PPIA (Peptidylprolyl Isomerase A) was used as a reference (Taqman Gene Expression Assays Rh02832197_gH, PPIA; Applied Biosystems PN4351370). The amount of viral RNA relative to PPIA mRNA was expressed by tabulating the difference in Ct values for each sample.

### Determination of neutralizing antibody titers

SARS-CoV-2 reporter viral particle neutralization (RVPN) was performed using the Wuhan-Hu-1 spike sequence (GenBank: MN908947.3) modified by addition of the D614G mutation and removal of 21 C-terminal amino acids demonstrated to enhance incorporation into viral particles. Recombinant vesicular stomatitis virus (VSV) containing firefly luciferase gene (Kerafast, Boston, MA) and incorporating SARS-CoV-2 spike were added to heat-inactivated samples diluted four-fold, together with positive, negative and no-serum controls. The resulting mix was incubated and then added to 96-well plates containing ACE2 and TMPRSS2 expressing HEK293T cells. Eighteen to 24 h later, luciferase activity was measured on a chemiluminescence reader (BMG CLARIOStar, BMG LABTECH Inc., Cary, NC) after lysing the cells. Neutralization titers were calculated as a percentage of no-serum control and the NT50 was estimated from the dilution curve using Prism8 (GraphPad Software, San Diego, CA). Titers below 40 were considered non-neutralizing.

### Measurement of virus-specific binding antibodies

The plasma donor pools and plasma samples from all treated animals were also tested on 2 additional assays that measure binding antibodies. The VITROS^®^ Anti-SARS-CoV-2 Total Test (which measures total antibodies to the SARS-CoV-2 virus (IgG, IgM, IgA and other isotypes) was performed following the manufacturer’s instructions [46]. Briefly, serum samples are quickly vortexed, loaded on Ortho VITROS XT-7200 or 3600 instruments (Ortho Clinical Diagnostics, Raritan, NJ) and programmed for the CoV2T test following the manufacturer’s instructions. The S1 antigens coated on the assay wells bind S1 antibodies from human serum which, in turn, bind to a secondary HRP-labeled S1 antigen in the conjugate reagent forming a sandwich. The addition of signal reagent containing luminol generates a chemiluminescence reaction that is measured by the system and quantified as the ratio of the signal relative to the cut-off value generated during calibration. In addition, the donor plasma pools and samples from the treated animals were also assessed for antibody responses to SARS-CoV-2, SARS-CoV, middle east respiratory syndrome coronavirus (MERS-CoV), various seasonal human coronaviruses (HCoV-229E, HCoV-NL63, HCoV-HKU1, and HCoV-OC43), influenza, and several other common-cold viruses by using a coronavirus antigen microarray (CoVAM) described earlier [18, 47].

### Measurement of cytokines and chemokines in plasma

Plasma cytokines and chemokines were measured using the Cytokine 29-Plex Monkey Panel (Invitrogen™) which is a multiplex microbead fluorescent assay utilizing the Luminex™ platform. The assay was run according to manufacturer’s instructions.

### Serum biochemistry

Biochemistry analysis on serum samples was performed using Piccolo® BioChemistry Plus disks, that were run on the Piccolo® Xpress Chemistry Analyzer (Abbott), according to the manufacturer’s instructions. This panel includes alanine aminotransferase (ALT), albumin, alkaline phosphatase (ALP), amylase, aspartate aminotransferase (AST), C-reactive protein, calcium, creatinine, gamma glutamyltransferase (GGT), glucose, total protein, blood urea nitrogen (BUN), and uric acid.

### Measurement of Innate and adaptive immune responses by Flow cytometry

The Innate and adaptive immune responses were analyzed at days 0, 1, 3, 5, and 7. Fresh whole blood (150µl) from each study animal was stained with a panel of fluorophore conjugated monoclonal antibodies against the following surface antigens: CD3, CD4, CD8, CD11c, CD14, CD16, CD20, CD28, CD66, CD95, CD123, HLA-DR, and PD1 for 30 minutes at room temperature. Cells were subsequently lysed with 1000µL of FACS lyse (BD Biosciences, USA) then washed twice with 1X FACS buffer. Cells were fixed and the permeabilized with 100µL of FoxP3 fixation (BD Biosciences, USA) for 10 minutes in the dark at room temperature, followed by washing with 1X FoxP3 wash buffer. Intracellular staining was performed with Ki-67 for 45 minutes at 4°C. Fluorescence was measured on the same day using a BD FACSymphony with FACS Diva version 8.0.1 software. Compensation, gating, and analysis were performed using FlowJo (Versions 9 and 10). Antibody/reagent details are given in table S5.

### Euthanasia and evaluation of pathology

All animals were euthanized at day 7 after infection with an overdose of pentobarbital and subjected to a full necropsy under BSL-3 conditions. The lungs were harvested and each lobe separated. All 7 lung lobes were cannulated with 18-gauge blunt needles. A small peripheral section of the cannulated lobe was clamped off and removed to be saved for possible viral RNA analysis. Then the lung lobes were slowly infused with neutral buffered formalin at 30 cm fluid pressure. Once fully inflated (approx. 30 mins) the main bronchus was tied off and the lungs were placed in individual jars of formalin and fixed for 72 hours. Then they were sliced from the hilus towards the periphery into slabs approximately 5mm thick. Each slab was placed into a cassette recording its position in the stack and with further division of the slab into smaller pieces if required to fit into the cassette. Tissues were then held in 70% ethanol until processing. A full set of remaining tissues was harvested at necropsy, trimmed into cassettes and then fixed in 10% neutral buffered formalin for 72 hours before transfer into 70% ethanol. The lung and all other tissues were sent for routine tissue processing and paraffin embedding followed by sectioning at 5 µm and generation of H&E stained slides.

Slides from every other slab of all 7 lung lobes were examined independently by 2 ACVP board certified pathologists. Using a 1.5 mm spaced grid, 25 randomly selected x40 fields from each slide were evaluated for the severity of the interstitial inflammation (mostly mononuclear cells sometimes with neutrophils). Each field was graded on a scale of 0-4, as described previously [27], based on the number of cells expanding the alveolar interstitium: grade 1, 1-2 cells thick; grade 2, 3-4 cells thick; grade 3, 5-6 cells thick; grade 4, >6 cells thick. The score is allocated according to the most severe region within the field. Altogether, this method resulted in evaluating between 710 to 1634 microscopic fields (at x40 magnification) per animal. Then a weighted-average overall score was calculated that takes into account the number of readings per lung lobe (i.e., larger lung lobes, having more readings, contribute more to the overall score than smaller lobes).

The other tissues were also examined by a pathologist but no other significant lesions were identified.

### Statistical analyses

Statistical analyses were performed using Prism version 9 (GraphPad), with selection of the test as outlined in the results. P values of <0.05 were considered statistically significant.

## Acknowledgment

We are grateful to David Bennett, Wilhelm Von Morgenland, Miles Christensen, Vanessa Bakula, James Schulte, Jose Montoya, Joshua Holbrook, John McAnelly, Sadie Rogers, Greg Hodge, Chelsea Schiano, and Nicole Drazenovich for expert technical assistance; Sheri Hild (ORIP, NIH) for assistance with funding acquisition; Que Dang (DAIDS/NIAID/NIH) for useful discussions; Clint Florence (DMID/NIAID/NIH) and BEI Resources for providing the virus.

Research reported in this publication was supported by the Office of Research Infrastructure Program, Office of The Director, National Institutes of Health under Award Number P51OD011107; NIAID grants 1R21AI143454-02S1 and 3RF1AG061001 to S. S. Iyer. The content is solely the responsibility of the authors and does not necessarily represent the official views of the National Institutes of Health. The funders had no role in study design, data collection and analysis, decision to publish, or preparation of the manuscript.

## SUPPLEMENT

**Figure S1:**
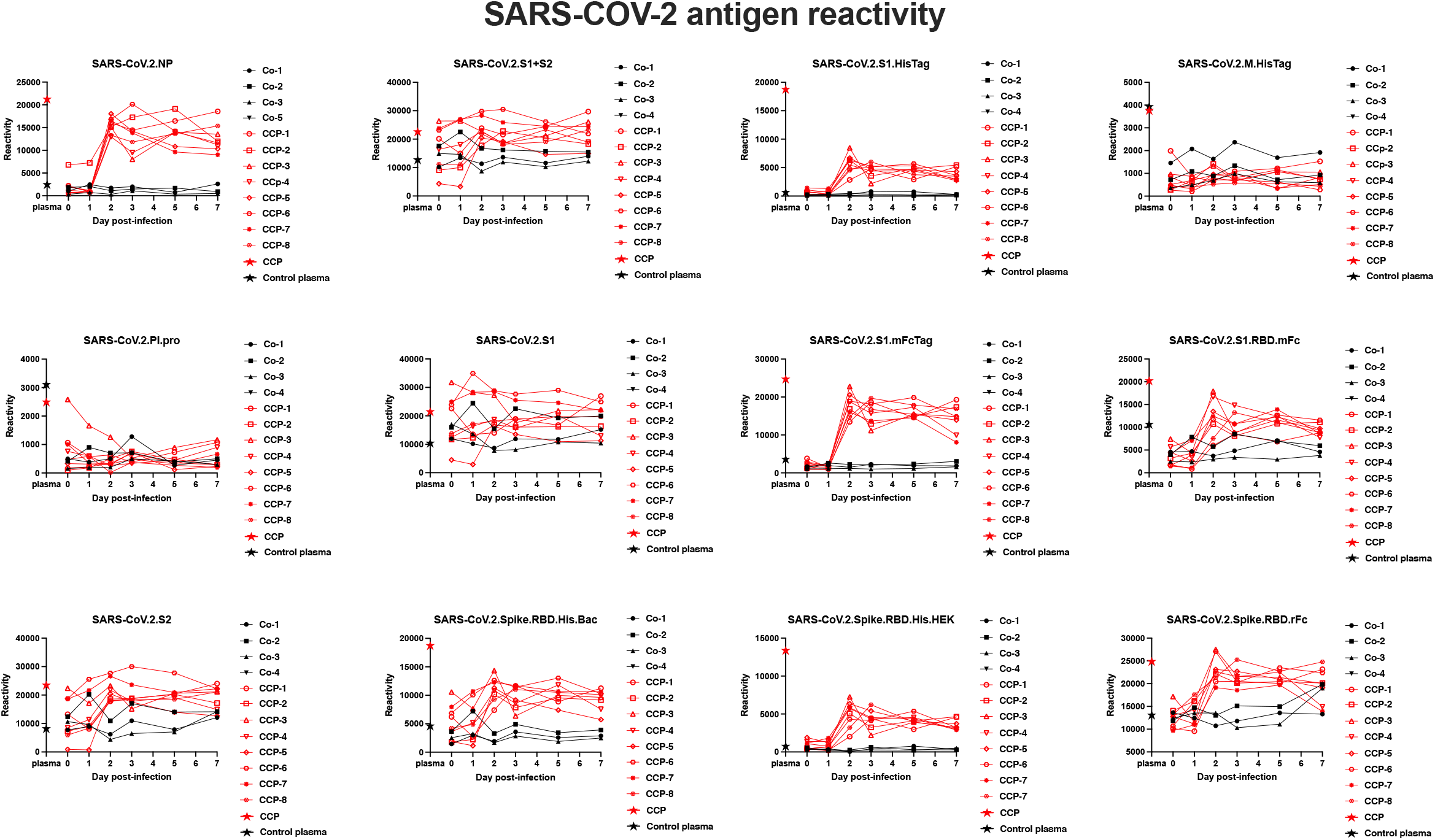
Reactivity of macaque plasma to SARS-CoV-2 antigens following human plasma infusion. Plasma collected of the macaques before and after infusion with pooled CCP or normal control plasma was tested by coronavirus antigen microarray assay (COVAM). The data on reactivity to SARS-CoV-2 antigens in this assay are represented as individual graphs. The reactivity of the CCP and normal plasma (see **Fig. 1A**) is indicated on the Y-axis as red and black stars, respectively, to demonstrate the dilution effect after transfusion into the macaques.

**Fig. S2.**
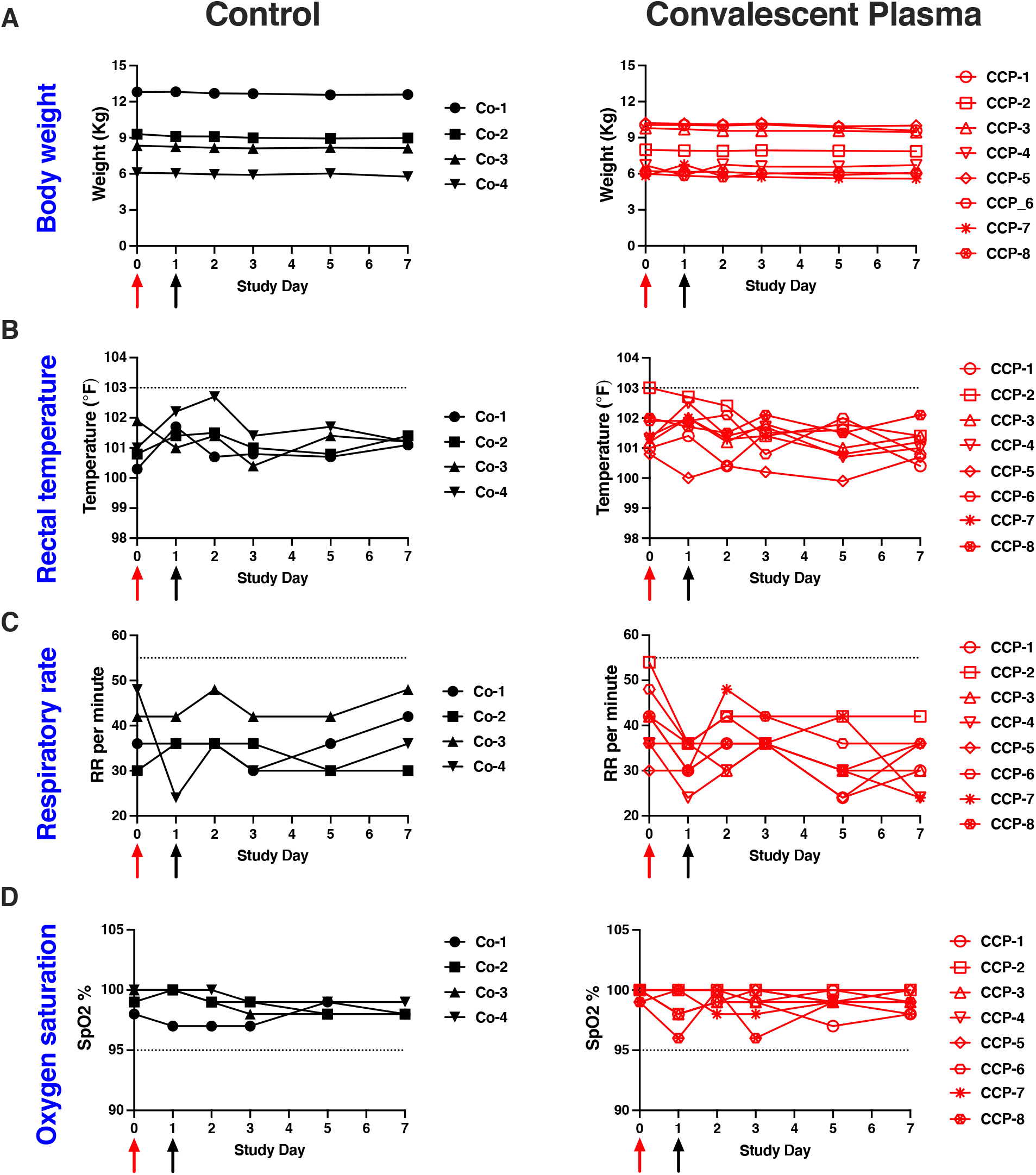
Clinical measurements collected at time of sedation. Red and black arrows indicate time of virus inoculation and monoclonal antibody administration on days 0 and 1, respectively. (A) Body weight remained stable. (B) Rectal temperature; horizontal line indicates the cut-off of 103° F, above which ketoprofen treatment was administered. (C) Respiratory rate; the horizontal line indicates a cut-off value of 55 (per minute) as upper normal range. (D) Oxygen saturation measured by pulse oximetry; the horizontal line indicates 95% as the lower end cut-off of the normal range. Total clinical scores, including the markers not graphed above, are presented in Fig. 4.

**Fig. S3.**
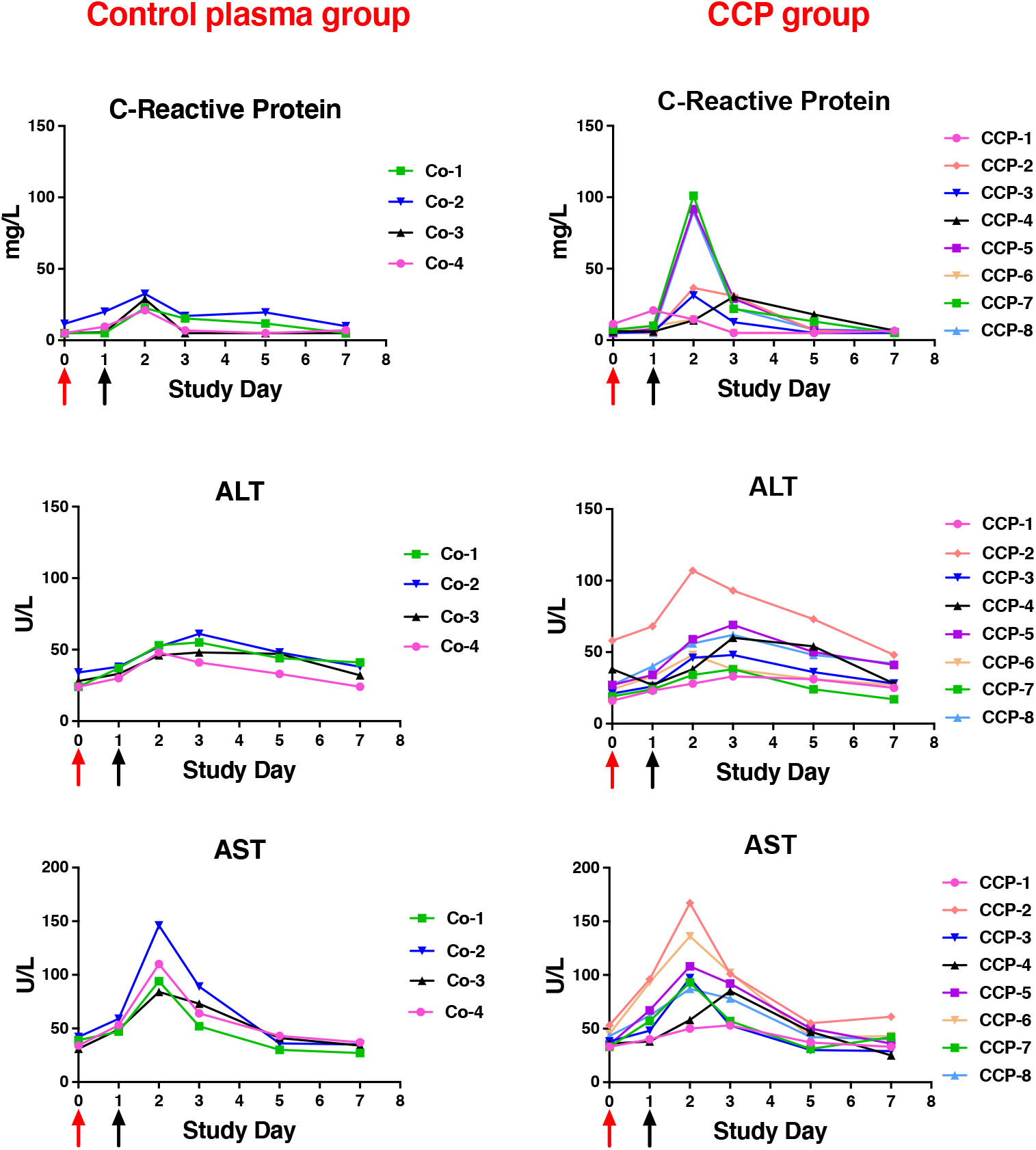
Time course of serum chemistry markers in SARS-CoV-2 inoculated animals. Biochemistry analysis on serum samples was performed using Piccolo® BioChemistry Plus disks. (A) through (C) present C-reactive protein (CRP), alanine aminotransferase (ALT), and aspartate aminotransferase (AST), which showed transient changes during the early stages of infection regardless of the study group. Other markers in the panel did not show any obvious changes. Red and black arrows indicate time of virus inoculation and plasma administration on days 0 and 1, respectively.

**Fig. S4.**
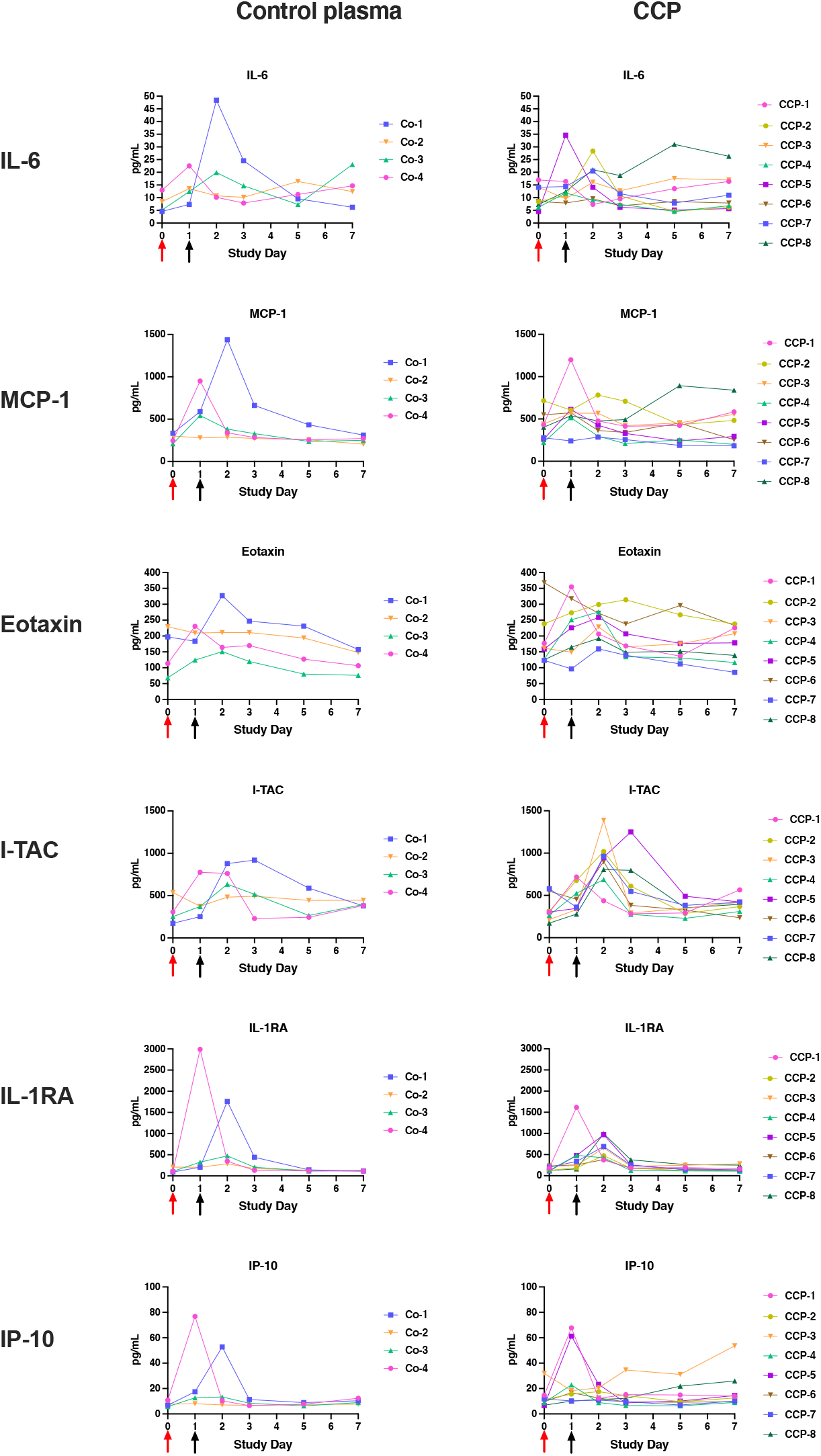
Time course of cytokines and chemokines in plasma of SARS-CoV-2 inoculated animals. Cytokines and chemokines were measured in plasma using established Luminex-based methodology (see methods section). Red and black arrows indicate time of virus inoculation and plasma administration on days 0 and 1, respectively. Markers on this figure represent ones that showed the most visible changes after infection. For other markers, see **Fig. S5**.

**Fig. S5.**
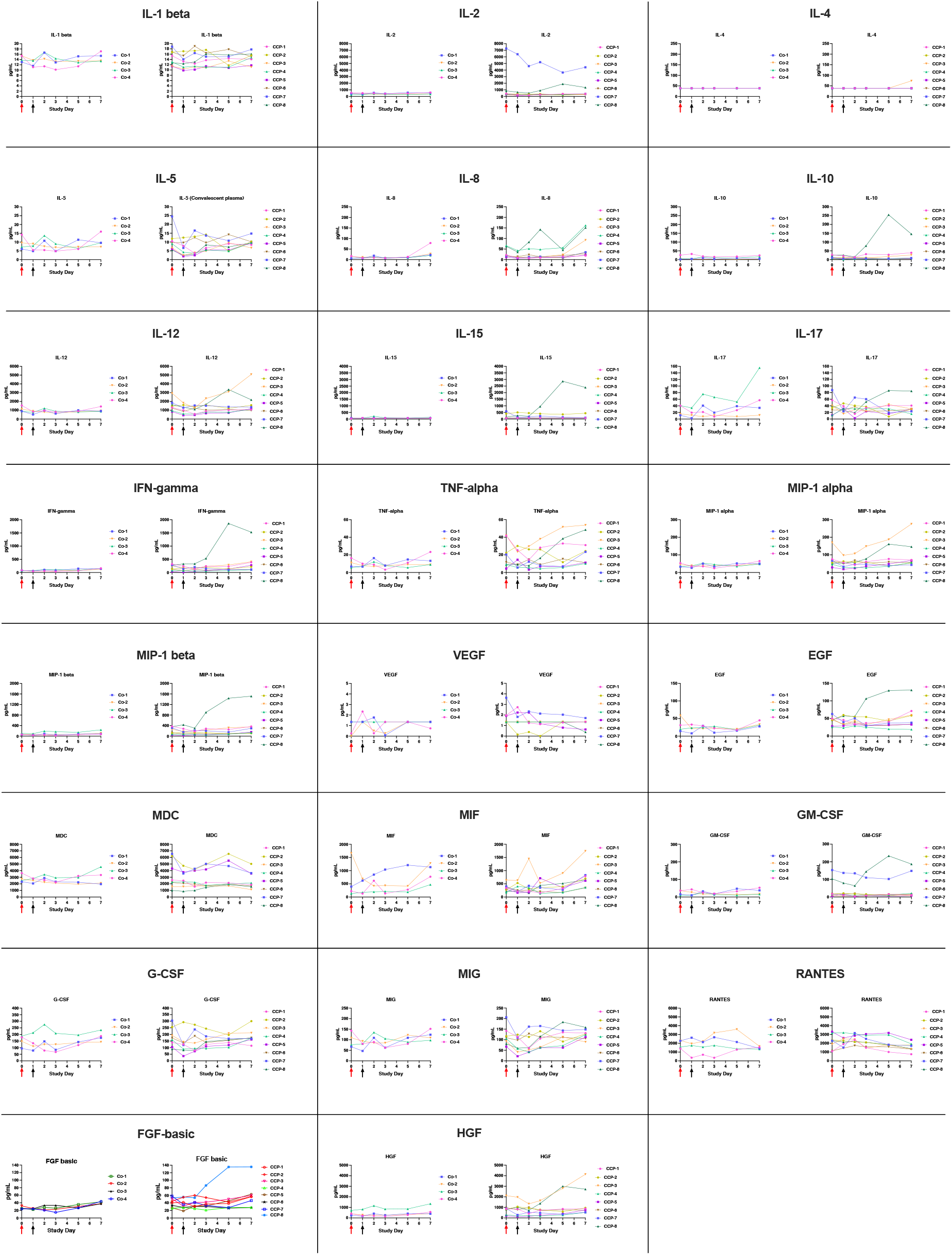
Time course of additional cytokines and chemokines in plasma of SARS-CoV-2 inoculated animals. Cytokines and chemokines presented in this panel were ones that did not show consistent changes among animals. The legend is the same as that of **Fig. S4**.

**Fig. S6.**
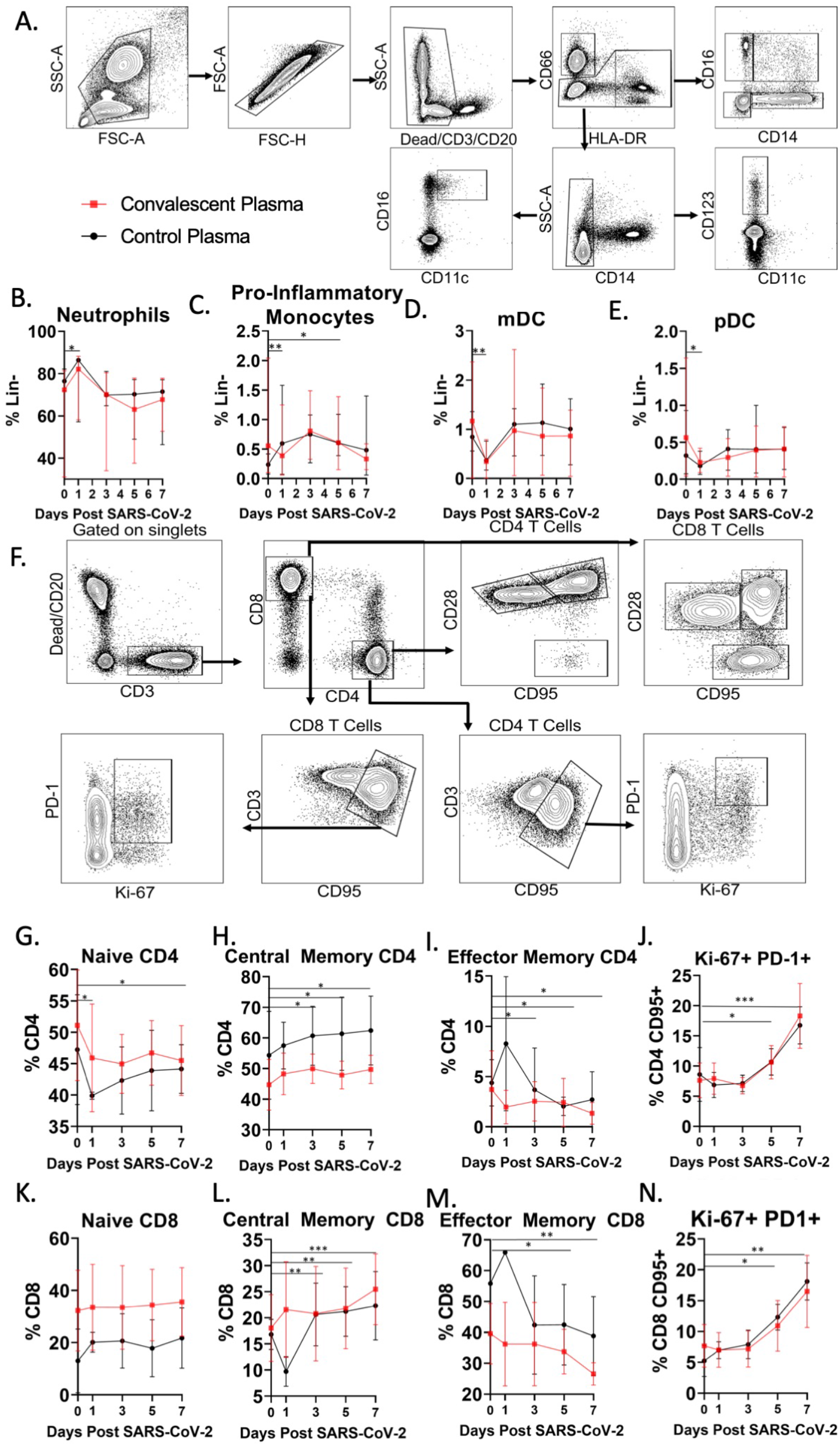
Innate and adaptive immune responses following infection. (A) Representative gating strategy for innate immune cells in whole blood. Fluorochromes used: CD66:APC, CD20/CD3/Dead: APC-Cy7,, Ki67:AF488, CD14:AF700, CD123:BV421, CD16:BV605, HLA-DR:BV786, CD11c:PE-Cy7. Kinetics of circulating neutrophils, proinflammatory monocytes, mDCs, and pDCs measured at 0,1,3,5, and 7 days post SARS-CoV-2 infection. (B) Representative T cell gating strategy from whole blood. Fluorochromes used: CD25: APC, CD20/Dead: APC-Cy7, Ki67:AF488, CD3:AF700, CD95:BUV737, CD8:BUV805, CD4:BV650, CD69:BV711, CD28:PECF594, PD-1:PE-Cy7. Kinetics of circulating naïve, central memory, effector memory populations, and Ki67+PD-1+ memory cells of CD4 T cell. Kinetics of circulating naïve, central memory, effector memory, and CD69+ effector memory CD8 T cells. Significance was calculated using one tailed paired t test comparing pooled convalescent plasma and normal plasma animals against Day 0 *p=0.05, **p=0.01, ***p=0.001. Statistical analysis yielded no significant different between convalescent and normal plasma groups.

**Fig. S7.**
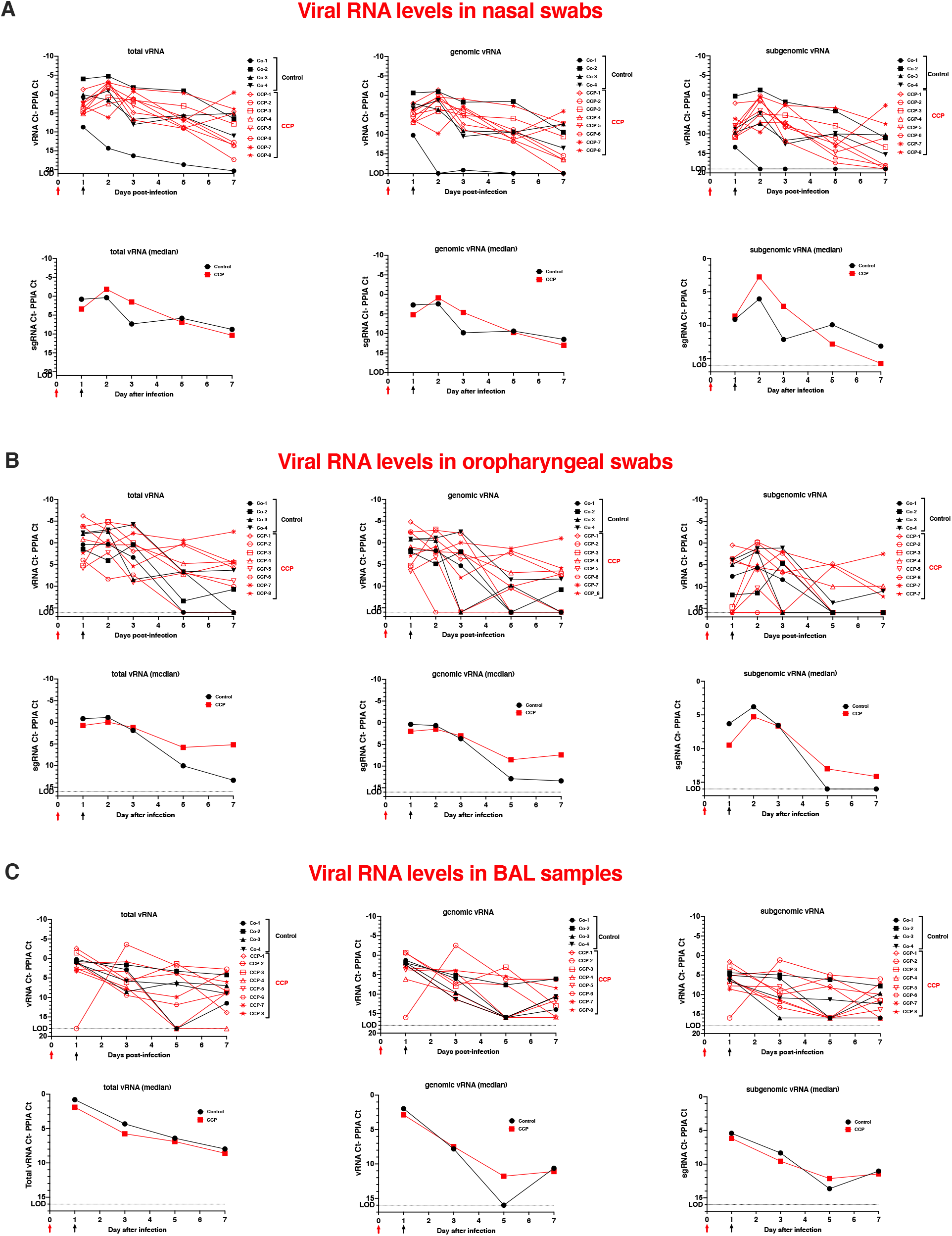
Viral RNA levels in nasal swabs, oropharyngeal swabs and BAL samples. Nasal swabs (A), oropharyngeal swabs (B) and BAL (cell pellets with supernatant) (C) were tested by RT-qPCR for total, genomic and subgenomic viral RNA, and the housekeeping gene PPIA mRNA. Viral RNA levels are expressed relative to PPIA mRNA by graphing the difference in Ct values. For each sample type, the top figures show the individual data (with the intersection of X-axis and Y-axis set near the limit of detection); the bottom figures display the median values per group. Red and black arrows indicate time of virus inoculation and monoclonal antibody administration on days 0 and 1, respectively.

**Fig. S8.**
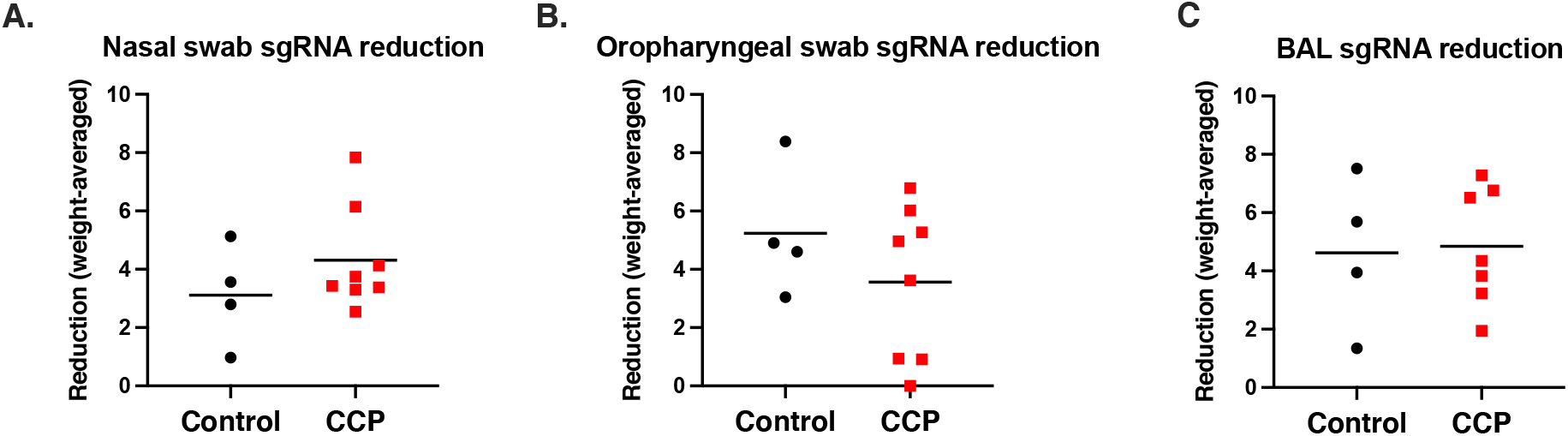
Lack of effect of convalescent plasma on sgRNA kinetics in nasal and oropharyngeal swabs and BAL of SARS-CoV-2 infected macaques. A weighted average analysis was performed on the sgRNA data from nasal and oropharyngeal swabs and BAL (**Fig. S7**) to calculate the relative decline of viral RNA (relative to cellular mRNA in the sample) from day 1 to day 7. For each animal, the AUC of relative sgRNA per cellular mRNA over time was tabulated using day 1 as baseline value, and then divided by 6 days to get the weighted average in the decline of sgRNA over the 6-day time period. Lines indicate mean values. On panel C, animal CCP-2 was excluded, as it had no detectable viral RNA in the BAL sample, which precluded this analysis. Statistical analysis revealed no effects between the control and CCP groups (panel A, p=0.29; panel B: p=0.30; panel C, p=0.88; unpaired t-test).

**Fig. S9.**
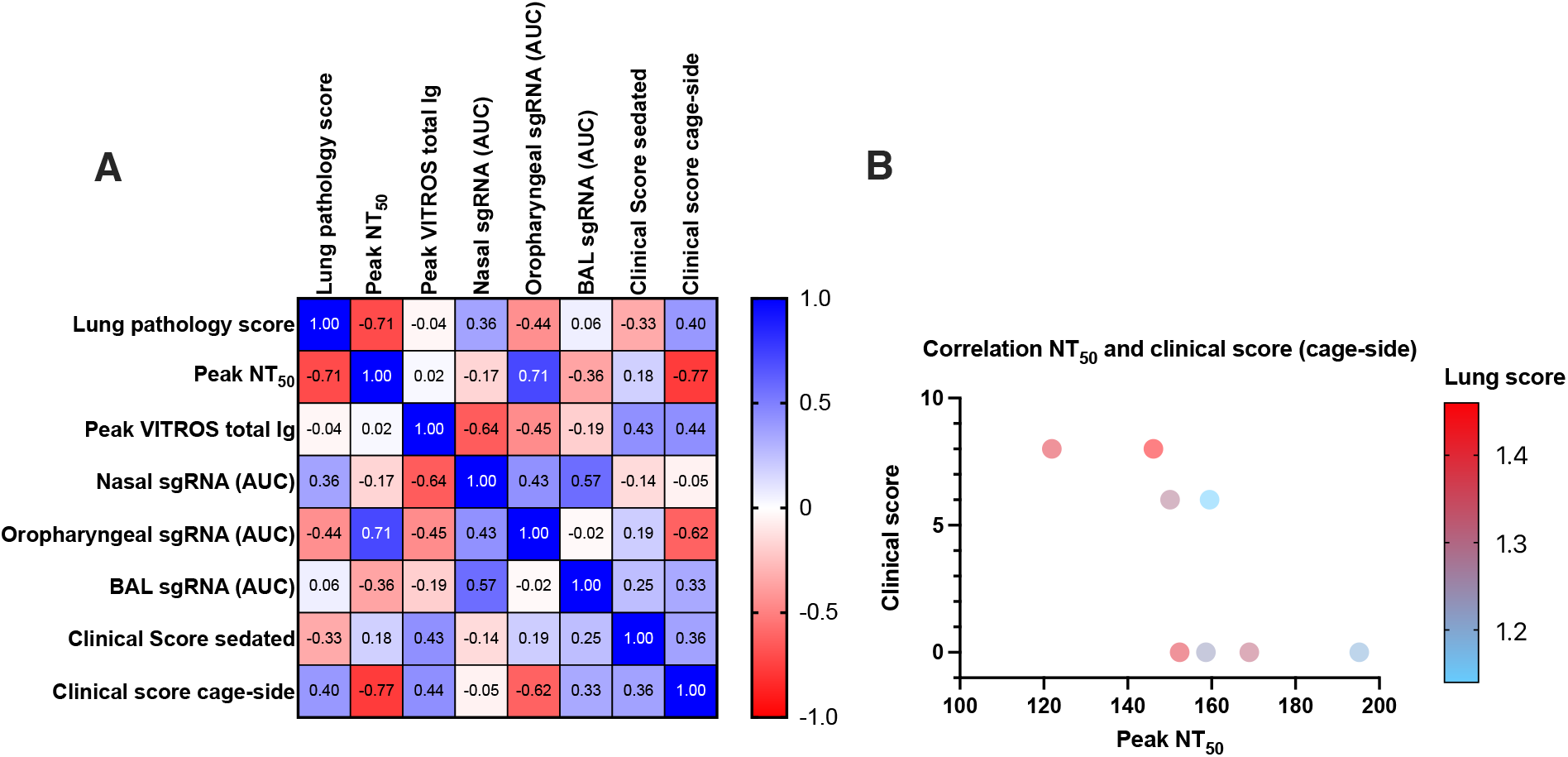
Multivariable correlation analysis on CCP-treated animals. Multivariate analysis was performed on the 8 CCP-treated animals only. (A). Spearman r correlation matrix in heatmap format. For this analysis, the markers used are the same ones as in Fig. 7. (B) Correlation between neutralizing antibody peak NT_50_ values and clinical scores based on cage-side observations (Spearman r = −0.77; p=0.04). The labels next to each symbol indicate the individual animals.

**Fig. S10:**
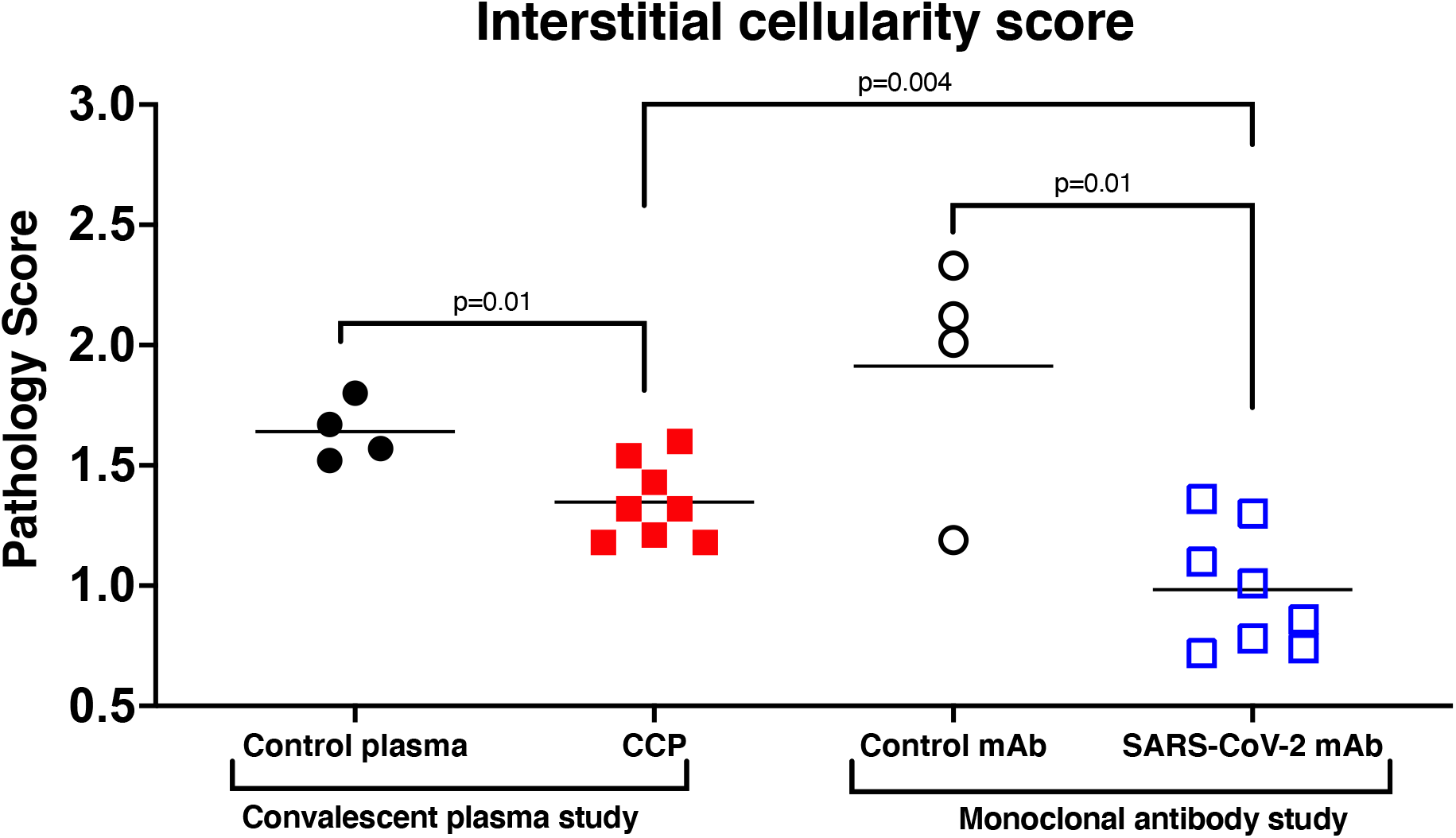
Comparison of effects of convalescent plasma and monoclonal antibodies on reducing lung inflammation in SARS-CoV-2 infected macaques. In an earlier study that used the same experimental procedures, we demonstrated that a combination of 2 potent anti-SARS-CoV-2 monoclonal antibodies (mAb; C135-LS and C144-LS) administered one day after virus inoculation reduced lung interstitial cellularity scores in 8 treated animals comparison to 4 animals treated with a control mAb [27]. Comparison of the CCP-treated and SARS-CoV-2 mAb-treated groups demonstrated that mAbs are more effective than CCP in reducing interstitial cellularity scores (p=0.004, unpaired t test). Because in the monoclonal antibody study, scores were based on 3 lung lobes, the data of the CCP study presented in this figure are tabulated based on those same 3 lung lobes; using the data of all 7 lung lobes on the current CCP study (presented in **Fig. 5**) resulted in the same conclusions.

**Table S1:** Preparation of pooled COVID convalescent plasma. Due to limited amount available of the plasma with the highest titer (W084520000915), a maximal amount of this highest-titer plasma was used and mixed with the 2 other units at a ratio of 60:20:20 in order to administer the maximum absolute amount of convalescent plasma-derived neutralizing antibodies to the animals. ^1^NT_50_ and NT_80_ titers determined by RVPN assay. ^2^Top percentile values of individual plasma units and the pooled CCP are based on 223 convalescent plasma samples with median NT_50_ titer of 784. ^3^Signal to cut-off ratio values on VITROS^®^ Total Ig assay (which measures anti-spike IgG, IgM and IgA). A value of ≥1 is considered reactive.

| **Truncated DIN** | **% of total pool** | **NT<sub>50</sub><sup>1</sup>** | **Top percentile based on NT<sub>50</sub><sup>2</sup>** | **NT<sub>80</sub><sup>1</sup>** | **VITROS S/CO<sup>3</sup>** |
| --- | --- | --- | --- | --- | --- |
| W084520000915 | 60% | 18,922 | 1% | 2,313 | 736 |
| W041020069696 | 20% | 1,135 | 40% | 541 | 345 |
| W041120015179 | 20% | 1,350 | 35% | 454 | 506 |
| **Pooled CCP** | **100%** | **3,003** | **20%** | **1,113** | **684** |

**Table S2.**
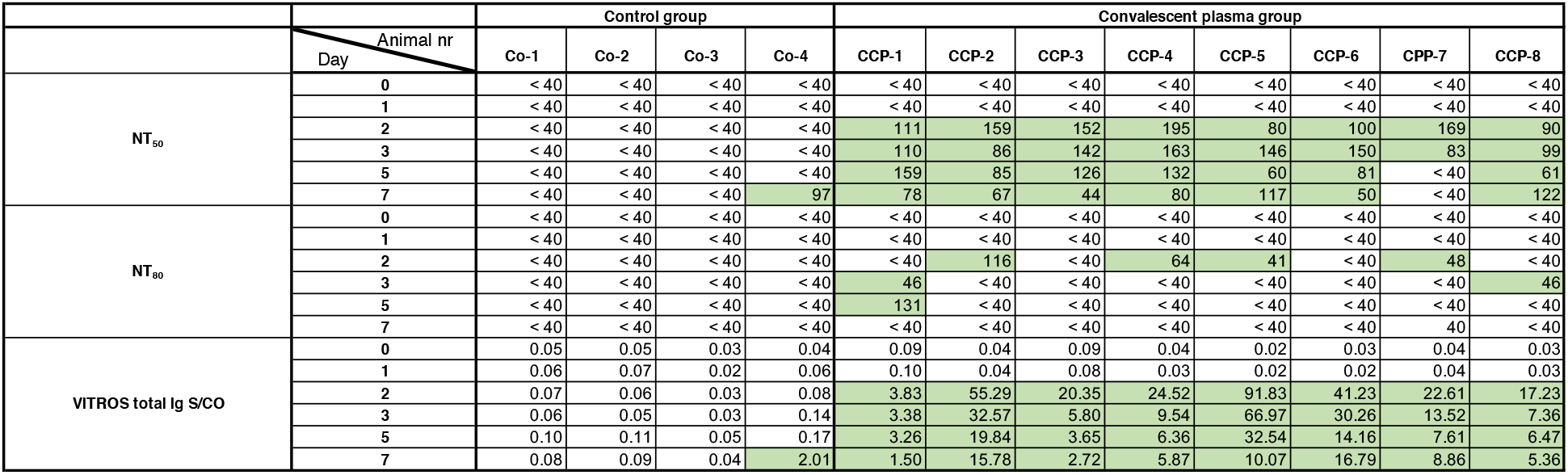
SARS-CoV-2 neutralizing and anti-spike antibodies in serum of macaques. Animals were inoculated with SARS-CoV-2 on day 0, and control or convalescent plasma was infused on day 1. 50% and 80% neutralization titers (NT_50_ and NT_80_) in serum were measured by a RVPN assay. Total Ig against spike protein was determined by the VITROS^®^ assay. Green shading indicates values above the cut-off of the respective assay.

**Table S3.** Summary of radiological scoring. All thorax radiographs were scored blinded by a veterinary radiologist, with scores of 0 to 3 assigned to each of the 7 lung lobes. For each time point, the total score of all lung lobes was tabulated. Thus, the maximum score per time point is 21.

| Control plasma |  |  | Convalescent plasma |  |  |
| --- | --- | --- | --- | --- | --- |
| Animal number | Day | Total Score | Animal number | Day | Total Score |
| Co-1 | 0 | 0 | CCP-1 | 0 | 0 |
|  | 1 | 0 |  | 1 | 0 |
|  | 3 | 1 |  | 3 | 0 |
|  | 5 | 1 |  | 5 | 0 |
|  | 7 | 2 |  | 7 | 0 |
| Co-2 | 0 | 0 | CCP-2 | 0 | 0 |
|  | 1 | 0 |  | 1 | 0 |
|  | 3 | 0 |  | 3 | 1 |
|  | 5 | 0 |  | 5 | 0 |
|  | 7 | 0 |  | 7 | 0 |
| Co-3 | 0 | 0 | CCP-3 | 0 | 0 |
|  | 1 | 0 |  | 1 | 0 |
|  | 3 | 1 |  | 3 | 1 |
|  | 5 | 0 |  | 5 | 0 |
|  | 7 | 0 |  | 7 | 1 |
| Co-4 | 0 | 0 | CCP-4 | 0 | 0 |
|  | 1 | 0 |  | 1 | 1 |
|  | 3 | 0 |  | 3 | 1 |
|  | 5 | 0 |  | 5 | 1 |
|  | 7 | 0 |  | 7 | 0 |
|  |  |  | CCP-5 | 0 | 0 |
|  |  |  |  | 1 | 0 |
|  |  |  |  | 3 | 0 |
|  |  |  |  | 5 | 0 |
|  |  |  |  | 7 | 0 |
|  |  |  | CCP-6 | 0 | 0 |
|  |  |  |  | 1 | 0 |
|  |  |  |  | 3 | 0 |
|  |  |  |  | 5 | 0 |
|  |  |  |  | 7 | 0 |
|  |  |  | CCP-7 | 0 | 0 |
|  |  |  |  | 1 | 0 |
|  |  |  |  | 3 | 0 |
|  |  |  |  | 5 | 0 |
|  |  |  |  | 7 | 0 |
|  |  |  | CCP-8 | 0 | 0 |
|  |  |  |  | 1 | 0 |
|  |  |  |  | 3 | 2 |
|  |  |  |  | 5 | 1 |
|  |  |  |  | 7 | 0 |

**Table S4.** Animal demographics.

| **Group** | **Animal ID** | **Sex** | **Age at time of inoculation (months)** | **Body weight at time of inoculation (kg)** |
| --- | --- | --- | --- | --- |
| Control plasma | Co-1 | M | 195 | 12.81 |
| " | Co-2 | F | 169 | 9.31 |
| " | Co-3 | M | 167 | 8.34 |
| " | Co-4 | F | 110 | 6.09 |
| Convalescent plasma | CCP-1 | F | 196 | 10.05 |
| " | CCP-2 | M | 166 | 7.99 |
| " | CCP-3 | M | 163 | 9.79 |
| " | CCP-4 | F | 137 | 6.70 |
| " | CCP-5 | M | 135 | 10.22 |
| " | CCP-6 | F | 112 | 6.01 |
| " | CCP-7 | F | 111 | 5.82 |
| " | CCP-8 | M | 110 | 6.22 |

**Table S5.** Flow cytometry antibody and reagents.

| **S. No** | **Reagents** | **Source** | **Identifier** |
| --- | --- | --- | --- |
| 1. | AF488 anti-human Ki-67 (Clone B56) | BD Biosciences | Cat#558616 |
| 2. | AF700 anti-human CD14 (Clone MSE2) | BD Biosciences | Cat# 301822 |
| 3. | AF700 anti-human CD3 (Clone SP34-2) | BD Biosciences | Cat# 557917 |
| 4. | APC anti-human CD66 (Clone TET2) | Miltenyi Biotec | Order#130-118-539 |
| 5. | APC-Cy7 anti-human CD3 (Clone SP34-2) | BD Biosciences | Cat#557757 |
| 6. | APC-Cy7 anti-human CD20 (Clone 2H7) | BioLegend | Cat#302314 |
| 7. | APC-Cy7 anti-human live/dead | invitrogen | Ref#L34976A |
| 8. | BV421 anti-human CD123 (Clone 7G3) | invitrogen | Ref#48-1238-42 |
| 9. | BV605 anti-human CD16 (Clone 3G8) | BioLegend | Cat#302040 |
| 10. | BV650 anti-human CD4 (Clone L200) | BD Biosciences | Cat# 563737 |
| 11. | BV786 anti-human HLA-DR (Clone L243) | BioLegend | Cat#307642 |
| 12. | BUV737 anti-human CD95 (Clone DX2) | BD Biosciences | Cat# 564710 |
| 13. | BUV805 anti-human CD8 (Clone SK1) | BD Biosciences | Cat#612889 |
| 14. | PECF594 anti-human CD28 (Clone CD28.2) | BioLegend | Cat# 302942 |
| 15. | PECy7 anti-human CD11c (Clone 3.9) | invitrogen | Ref#25-0116-42 |
| 16. | PECy7 anti-human PD1 (Clone EH12.2H8) | BioLegend | Cat# 329918 |
| 17. | FACS lyse | BD Biosciences | Cat#349202 |
| 18. | FoxP3/ Transcription Factor Staining Buffer set | invitrogen | Cat#00-5523 |
| 19. | Brilliant stain buffer | BD Biosciences | Cat#563794 |

## Notes

### Competing Interest Statement

The authors have declared no competing interest.

